# The bivalent epigenetic regulator ULTRAPETALA1 promotes the Arabidopsis floral transition via recruitment of Polycomb histone methyltransferases for H3K27me3 deposition at the *FLC* locus

**DOI:** 10.1101/2025.05.06.652514

**Authors:** Qian Xing, Elena Shemyakina, Jingjing Tian, Vivien I. Strotmann, Alia Anwar, Xianglian Li, Ying Xiong, Tianyue Yang, Yucai Zheng, Li Pu, Yvonne Stahl, Ralf Müller-Xing, Jennifer C Fletcher

**Affiliations:** Jiangxi Provincial Key Laboratory of Plant Germplasm Innovation and Genetic Improvement, Lushan Botanical Garden, Chinese Academy of Sciences, Jiujiang 332900, China; Plant Epigenetics and Development, Lushan Botanical Garden, Chinese Academy of Sciences, Nanchang, 330114, China; College of Life Science, Nanchang University, Nanchang 330047, China; Plant Gene Expression Center, United States Department of Agriculture - Agricultural Research Service, Albany, CA 94710, USA; Department of Plant and Microbial Biology, University of California, Berkeley, Berkeley, CA 94720, USA; Institute for Developmental Genetics, Heinrich-Heine University, Düsseldorf 40225, Germany; Institute for Plant Developmental Genetics, Goethe-University, Frankfurt 60438, Germany; Biotechnology Research Institute, Chinese Academy of Agricultural Sciences, Beijing 100081, China

**Keywords:** Arabidopsis, epigenetics, histone modification, FLC, flowering, methylation, PcG, ULT1

## Abstract

Floral induction is a major transition during the plant life cycle that contributes to reproductive fitness in annual and perennial plants. Flowering occurs in response to multiple environmental and endogenous cues. In *Arabidopsis thaliana*, many of these cues converge on the regulation of the *FLOWERING LOCUS C (FLC)* gene, which encodes a MADS domain transcription factor that functions as the main floral repressor. Here we demonstrate that the bivalent epigenetic regulator ULTRAPETALA1 (ULT1) is both necessary and sufficient to promote flowering. We identify ULT1 as a novel component of the autonomous pathway (AP) and show that it accelerates flowering by directly repressing *FLC* transcription. We demonstrate that ULT1 physically associates with the SWINGER (SWN) and CURLY LEAF (CLF) histone methyltransferase components of Polycomb Repressive Complex 2 (PRC2) to regulate *FLC* transcription by altering the accumulation of H3K27me3 marks. Our data indicate that ULT1 promotes the floral transition by interacting with SWN and CLF to recruit PRC2 to the *FLC* locus to deposit repressive histone methylation and reduce transcription of this key floral repressor.

## INTRODUCTION

In agriculture, flowering time is an important quantitative trait that is critical for pollination as well as yield of crop plants (Jung and Muller 2009). The floral transition represents a major developmental milestone in the plant life cycle that coordinates its entry into reproductive growth with seasonal environmental conditions. In response to a combination of environmental and endogenous cues, a dynamic cell fate switch occurs that directs the shoot apical meristem (SAM) to transition from generating vegetative organs such as leaves to floral meristems that produce the reproductive organs. Photoperiod (day length), vernalization (cold exposure), and ambient temperature are major environmental factors that regulate the floral transition, whereas endogenous cues come from chronological age, the phytohormone gibberellic acid (GA), and the autonomous pathway (AP). These various inputs are then integrated by a small number of floral regulatory factors to ensure that flowering occurs under the optimal conditions to maximize reproductive capacity.

The importance of this developmental switch is reflected by the fact that the floral transition is controlled by both transcriptional and epigenetic regulatory factors. The former are canonical transcription factors, whereas the latter are trithorax group (trxG) and Polycomb group (PcG) complexes that confer stable gene activation and repression states, respectively. The PcG complex Polycomb Repressive Complex 2 (PRC2) was first identified in *Drosophila melanogaster* and is required for maintaining the correct spatiotemporal expression of many genes during animal and plant development (Margueron and Reinberg 2011). PRC2 catalyzes histone 3 lysine 27 tri-methylation (H3K27me3), a histone modification associated with chromatin condensation and transcriptional repression (Holoch and Margueron 2017). Drosophila PRC2 consists of four core components that are conserved in plants. Arabidopsis contains three E(z) homologous genes, *CURLY LEAF (CLF), SWINGER (SWN)* and *MEDEA (MEA)*, that encode histone methyltransferase catalytic units (Goodrich et al. 1997; Grossniklaus et al. 1998; Chitvivattana et al. 2004). It also contains three Su(z)12 homologous genes, *FERTILIZATION INDEPENDENT SEED 2* (*FIS2*), *EMBRYONIC FLOWER 2 (EMF2)* and *VERNALIZATION2* (*VRN2*) (Luo et al. 1999; Chitvivattana et al. 2004). *MEA* and *FIS2* are specifically involved in seed development (Grossniklaus et al. 1998), whereas *CLF* and *SWN* have partially redundant functions during postembryonic development, as do *EMF2* and *VRN2* (Goodrich et al. 1997; Chitvivattana et al. 2004). The PRC2 core protein ESC is encoded by a single copy gene *FERTILISATION INDEPENDENT ENDOSPERM* (*FIE*) that is essential for plant development (Ohad et al. 1996; Ohad et al. 1999). Finally, a small family of five *MSI*-like genes (*MSI1-5*, homologues of yeast *multicopy suppressor of IRA1*) show similarity to *Drosophila* p55, but only MSI1 and FVE (MSI4) are known to associate with other PRC2 components (De Lucia et al. 2008; Pazhouhandeh et al. 2011). At least two distinct PRC2 complexes, one including the subunits CLF, EMF2 and FIE and the other including SWN, VRN2 and FIE, repress the expression of *FLC* and its homologues *MAF4* and *MAF5* during vegetative development to promote flowering (De Lucia et al. 2008; Jiang et al. 2008; Sheldon et al. 2009). CLF also represses *FT* expression in seedlings (Jiang et al. 2008), illustrating a complex role for PRC2 in regulating the floral transition.

PRC2 complexes lack intrinsic DNA binding specificity and are recruited to their target genes by transcription factors that bind to short genomic *cis*-regulatory regions called Polycomb response elements (PREs). First identified in Drosophila (Kassis and Brown 2013; Simon and Kingston 2013), a half-dozen PRE *cis*-motif sequences have been characterized in Arabidopsis, including teloboxes and GA repeats that are considered necessary and sufficient for PRE activity (Deng et al. 2013; Xiao et al. 2017). Such *cis*-regulatory elements are bound by members of the C2H2 zinc finger family, the APETALA2-like family, and the BASIC PENTACYSTEINE family of transcription factors, among others (Xiao et al. 2017).

Both transcriptional and epigenetic factors are critical for the function of the AP, an endogenous regulatory activity that acts independently of environmental sensing. Mutants of the AP show delayed flowering in both long day and short days conditions, whereas their flowering can be accelerated by vernalization (Koornneef et al. 1991; Koornneef et al. 1998; Wu et al. 2020; Kyung et al. 2022). The AP consists of a heterogenous group of proteins with functions in chromatin modification, RNA processing, spliceosome activity and transcription elongation (Wu et al. 2020; Kyung et al. 2022). The activity of these proteins converges on the regulation of the MADS box transcription factor gene *FLOWERING LOCUS C (FLC)*, which encodes the central floral repressor in *Arabidopsis thaliana*. Following germination, high *FLC* expression levels are maintained by elevated H3K4me3 and H3K36me3 chromatin marks, mediated by COMPASS-like trxG complexes (Jiang et al. 2011) and EARLY FLOWERING IN SHORT DAYS (EFS/SDG8) (Zhao et al. 2005; Xu et al. 2008), that confer an open chromatin configuration at the locus and suppress premature flowering. The AP promotes the floral transition through the gradual repression of *FLC* transcription during vegetative growth (Kyung et al. 2022). One subset of AP components, FLD and FVE/MSI4, confer a repressive chromatin configuration at the *FLC* locus via histone demethylation and deacetylation, respectively (Liu et al. 2007; Yu et al. 2011; Yu et al. 2016). Another subset of RNA 3’-end processing factors such as FCA, FPA and FY promote the proximal polyadenylation of *FLC* antisense long noncoding RNA transcripts collectively called *COOLAIR* (Hornyik et al. 2010; Liu et al. 2010). FCA binds nascent *COOLAIR* transcripts and directly interacts with the PRC2 component CLF to enrich H3K27me3 repressive chromatin marks at the *FLC* locus (Swiezewski et al. 2009; Csorba et al. 2014; Tian et al. 2019). AP-mediated epigenetic silencing of *FLC* transcription via both mechanisms prevents FLC protein from accumulating and repressing the expression of downstream target genes that promote the floral transition. However, whether additional factors play a role in autonomous pathway activity at the *FLC* locus is unknown.

FLC and its homologue SHORT VEGETATIVE PHASE (SVP) form a complex that directly represses the expression of the floral integrator genes *FLOWERING LOCUS T (FT)* and *SUPPRESSOR OF OVEREXPRESSION OF CONSTANS (SOC1)* (Li et al. 2008). Light quality and duration are perceived in the leaves of vegetative plants by photoreceptors that regulate the expression of the zinc finger transcription factor gene *CONSTANS (CO)* (Amasino 2010). Under inductive photoperiods the accumulation of CO protein leads to the direct activation of *FT* (An et al. 2004), which encodes a mobile signal called florigen that moves from the leaves to the SAM via phloem transport (Corbesier et al. 2007). Once in the SAM, FT protein physically associates with the bZIP transcription factor FLOWERING LOCUS D (FD) (Abe et al. 2005; Wigge et al. 2005). The FD-FT complex then activates *SOC1*, which also responds to the vernalization, age and GA pathways (Lee and Lee 2010), and these factors induce downstream floral meristem identity genes (Abe et al. 2005; Wigge et al. 2005; Yoo et al. 2005; Lui et al. 2008) that direct the cells on the flanks of the SAM to assume floral meristem identity.

A potential candidate factor to function in PRC2 recruitment to target loci during the flowering process is the ULTRAPETALA1 (ULT1) epigenetic transcriptional regulator. Arabidopsis *ULT1* and its paralog *ULT2* encode closely related proteins consisting of a DNA-binding SAND domain and a cysteine-rich B-box-like domain (Carles et al. 2005). The B-box-like domain of the rice OsULT1 protein plays a role in protein multimerization (Roy et al. 2019). Although the ULT proteins contain no known transcription activation or repression domains, based on the 3D structure of their DNA binding domain (DBD) they have been classified within a superclass of transcription factors whose DBDs consist of “alpha-helices exposed by beta-structures” (Blanc-Mathieu et al. 2024).

*ULT1* is expressed predominantly in Arabidopsis root, shoot and floral meristems as well as in developing reproductive organs (Carles et al. 2005; Ornelas-Ayala et al. 2020). ULT1 regulates hundreds of downstream target genes (Pu et al. 2013; Tyler et al. 2019) and functions in multiple plant developmental processes. In root apical meristems *ULT1* maintains the stem cell niche by regulating cell division rates in the quiescent center and preventing premature columella stem cell differentiation (Ornelas-Ayala et al. 2020). In shoot apical meristems *ULT1* acts partially redundantly with *ULT2* to restrict stem cell activity (Fletcher 2001; Monfared et al. 2013; Moreau et al. 2016). In reproductive meristems *ULT1* acts genetically as a trxG factor that promotes floral meristem termination by counteracting the activity of the PRC2 component CLF (Carles and Fletcher 2009), and later directs the patterning of the developing gynoecium (Monfared et al. 2013; Pires et al. 2014). Conversely, in young seedlings, ULT1 functions together with the PRC2 accessory component EMBRYONIC FLOWER 1 (EMF1) to prevent inappropriate seed gene expression after germination by binding seed gene loci and altering their histone methylation levels (Xu et al. 2018). The ability of the ULT1 protein to act cooperatively with PcG factors at some stages of development and antagonistically at others suggests a bimodal, stage-specific biochemical function for ULT1 and illustrates a key role for the gene during developmental transitions.

In this study we show that *ULT1* regulates another critical plant developmental transition, the transition from vegetative to reproductive development. We demonstrate that ULT1 promotes flowering under all photoperiods and is sufficient to induce flowering under both long day and short day conditions. We identify ULT1 as a novel component of the AP that indirectly promotes *FT* and *SOC1* expression through negative regulation of the *FLC* floral repressor MADS box gene. ULT1 directly binds to the *FLC* locus and represses its transcription by enhancing the deposition of repressive histone methylation marks and reducing the deposition of active marks. Our demonstration of a physical association between ULT1 and the PRC2 components SWN and CLF provides a molecular mechanism for how ULT1 promotes flowering, by binding to *FLC* and recruiting PRC2 to deposit repressive histone methylation marks at the locus and stably reduce the transcription of this key floral repressor gene.

## RESULTS

### *ULT1* but not *ULT2* is necessary and sufficient to promote flowering

Our previous work showed that mutations in the *ULT1* gene lead to slightly delayed bolting under constant light conditions (Carles et al. 2005). In addition, *ult1* alleles can rescue the early flowering phenotype of *LFYasEMF1* plants in which EMF1 activity is selectively eliminated from the leaf primordia, and can restore the expression of many mis-regulated genes in this background, including flowering time genes (Pu et al. 2013). Based on these data we hypothesized that *ULT1* might play a role in regulating the floral transition.

We investigated the roles of *ULT1* in regulating Arabidopsis flowering by characterizing the phenotypes of various *ult1* homozygous mutants in response to diverse flowering cues. The *ult1-2* allele is a cytosine to thymine nucleotide transition that replaces a conserved serine residue with a phenylalanine residue in the SAND domain (Carles et al. 2005), and behaves as a null allele with respect to shoot and floral meristem activity and gynoecium patterning (Monfared et al. 2013). The *ult1-3* allele is a T-DNA insertion 155 base pairs downstream of the start codon, early in the SAND domain region, and is a transcription null allele (Carles et al. 2005).

We observed delayed flowering phenotypes in both *ult1-2* and *ult1-3* plants when grown under long day (LD) conditions. Compared to wild-type Columbia-0 (Col-0) plants (Figure 1A), *ult1-2* (Figure 1B) and *ult1-3* (Figure 1C) plants flowered after producing an average of 5.27 and 9.4 additional total (rosette and cauline) leaves, respectively (Figure 1G), indicating that *ULT1* acts to accelerate flowering when the days are long. Although the *ULT2* gene is not expressed during vegetative development, it is transcribed in mature embryo shoot apical meristems (Carles et al. 2005). Therefore, we tested whether *ULT2* affected the floral transition using the *ult2-3* (Monfared et al. 2013) and *ult2-4* null alleles. The total number of leaves generated by *ult2-3* plants (Figure 1D) and *ult2-4* plants was not significantly different than that generated by Col plants (Figure 1G). Furthermore, the total leaf number of *ult1-2 ult2-3* (Figure 1E) and *ult1-3 ult2-4* (Figure 1F) double mutants was indistinguishable from that of *ult1* single mutants (Figure 1G). We conclude that *ULT1* but not *ULT2* is necessary to promote flowering under LD conditions.

**Figure 1.**
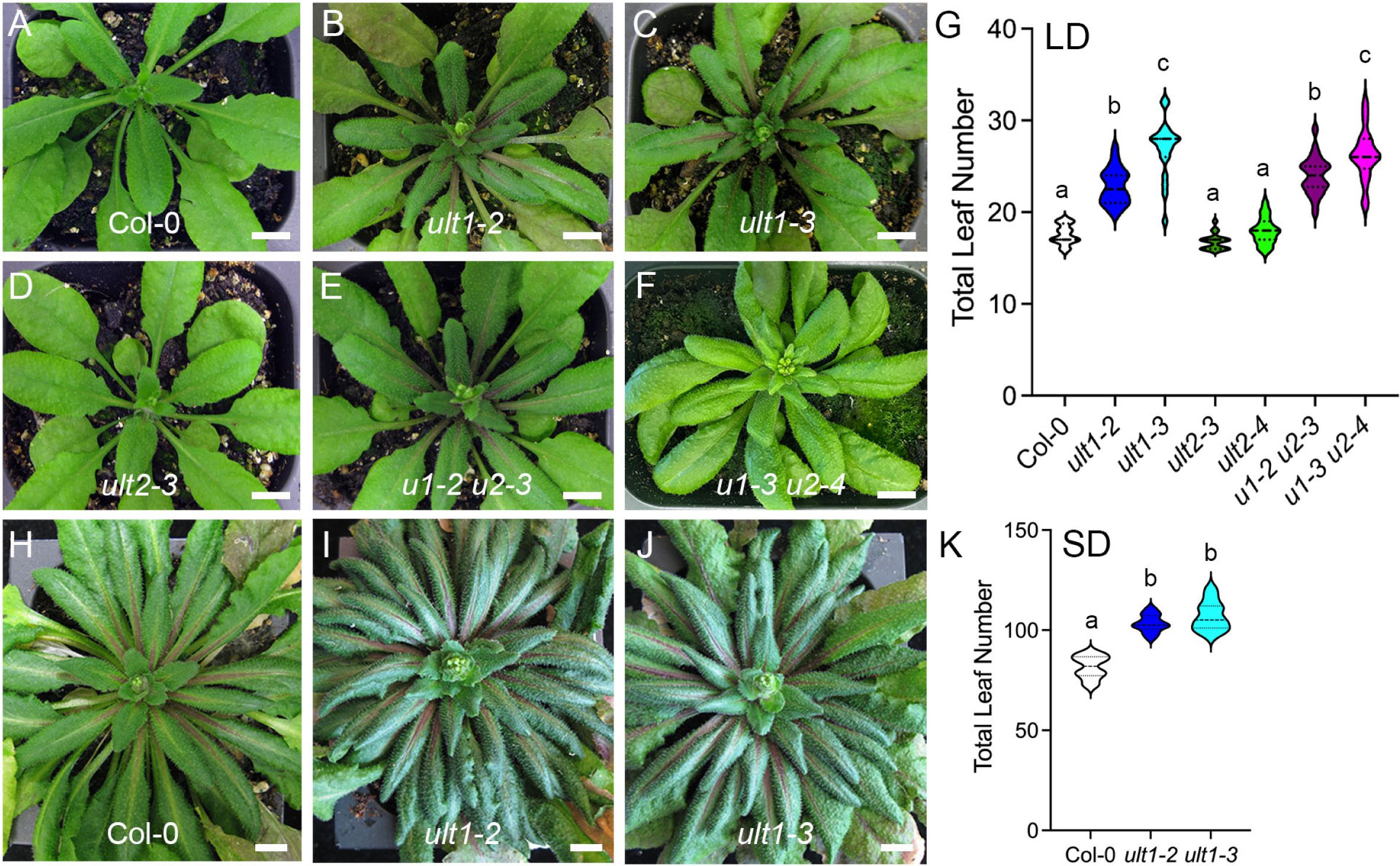
*ULT1* but not *ULT2* promotes flowering time under long day (LD) and short day (SD) conditions. **(A)** wild-type Col-0, **(B)** *ult1-2*, **(C)** *ult1-3*, **(D)** *ult2-3*, **(E)** *ult1-2 ult2-3* and **(F)** *ult1-3 ult2-4* plants grown to bolting under LD conditions. **(G)** Mean total leaf number under LD conditions. **(H)** wild-type Col-0, **(I)** *ult1-2* and **(J)** *ult1-3* plants grown to bolting under SD conditions. **(K)** Mean total leaf number under SD conditions. Lower case letters indicate statistically significant differences (P < 0.001) based on one-way ANOVA followed by Tukey’s test for multiple comparisons. n > 20. Scale bar, 1 cm.

Next, we tested whether *ULT1* promoted flowering under a short day (SD), 8-hour light photoperiod. We observed that the flowering of both *ult1-2* and *ult1-3* plants in SD conditions also was strongly delayed compared to that of Col plants (Figure 1H-K). These results demonstrate that *ULT1* promotes the transition to flowering independently of photoperiod.

We validated these results by analyzing the flowering phenotypes of *ult1-1* and *ult1-3* plants in the Landsberg *erecta* (L*er*-0) background. The *ult1-1* allele is a guanine to adenine nucleotide transition that replaces a conserved cysteine residue to a threonine residue at position 173 relative to the translational initiation site, causing semi-dominant phenotypes stronger than those of null alleles such as *ult1-3* (Carles et al. 2005). L*er* plants carry a weak allele of *FLC* conferred by a transposon insertion in the first intron of the gene (Gazzani et al. 2003), resulting in earlier flowering than other Arabidopsis accessions. *ult1-1* and *ult1-3* plants generated significantly more leaves than L*er* plants under LD conditions (Supplemental Figure 1). Furthermore, *ult1-1* and *ult1-3* plants generated significantly more leaves than L*er* plants under SD conditions (Supplemental Figure 1B). These results are consistent with the late flowering phenotype observed in the Col background under both light conditions.

Finally, we assessed the sufficiency of ULT1 to accelerate flowering in different photoperiods using two independent *35S*:*ULT1* L*er* over-expression lines. Under LD conditions plants from both *35S*:*ULT1* lines were significantly earlier flowering, making an average of two fewer leaves than L*er* plants (Supplemental Figure 2A-C), and showed additional leaf phenotypes caused by ectopic expression of floral homeotic genes in vegetative tissues (Carles and Fletcher 2009). Similarly, under SD conditions, *35S*:*ULT1* plants flowered after forming about half the number of leaves as L*er* plants (Supplemental Figure 2D-F). Because *ULT1* over-expression causes early flowering in both LD and SD conditions, we conclude that *ULT1* is sufficient to promote the floral transition.

### *ULT1* Acts in the Autonomous Pathway and Regulates Key Flowering Time Genes

Having established that *ULT1* promotes flowering independently of photoperiod, we analyzed whether it acts in other endogenous or environmental flowering time pathways. First, we analyzed whether *ULT1* acts in the endogenous gibberellic acid (GA) hormone pathway. GA signaling promotes flowering, most readily detectable under SD conditions, by increasing the expression of floral integrator genes (Srikanth and Schmid 2011). Compared to untreated plants (Figure 2A), wild-type Col plants grown under SD conditions showed accelerated flowering in the presence of exogenously added GA (Figure 2D and G). Similarly, both *ult1-2* and *ult1-3* plants grown under SD conditions showed accelerated flowering upon GA treatment (Figure 2E and 2F and 2G) compared to untreated plants (Figure 2B and 2C and 2G) Therefore, *ult1* mutants respond normally to GA treatment.

**Figure 2.**
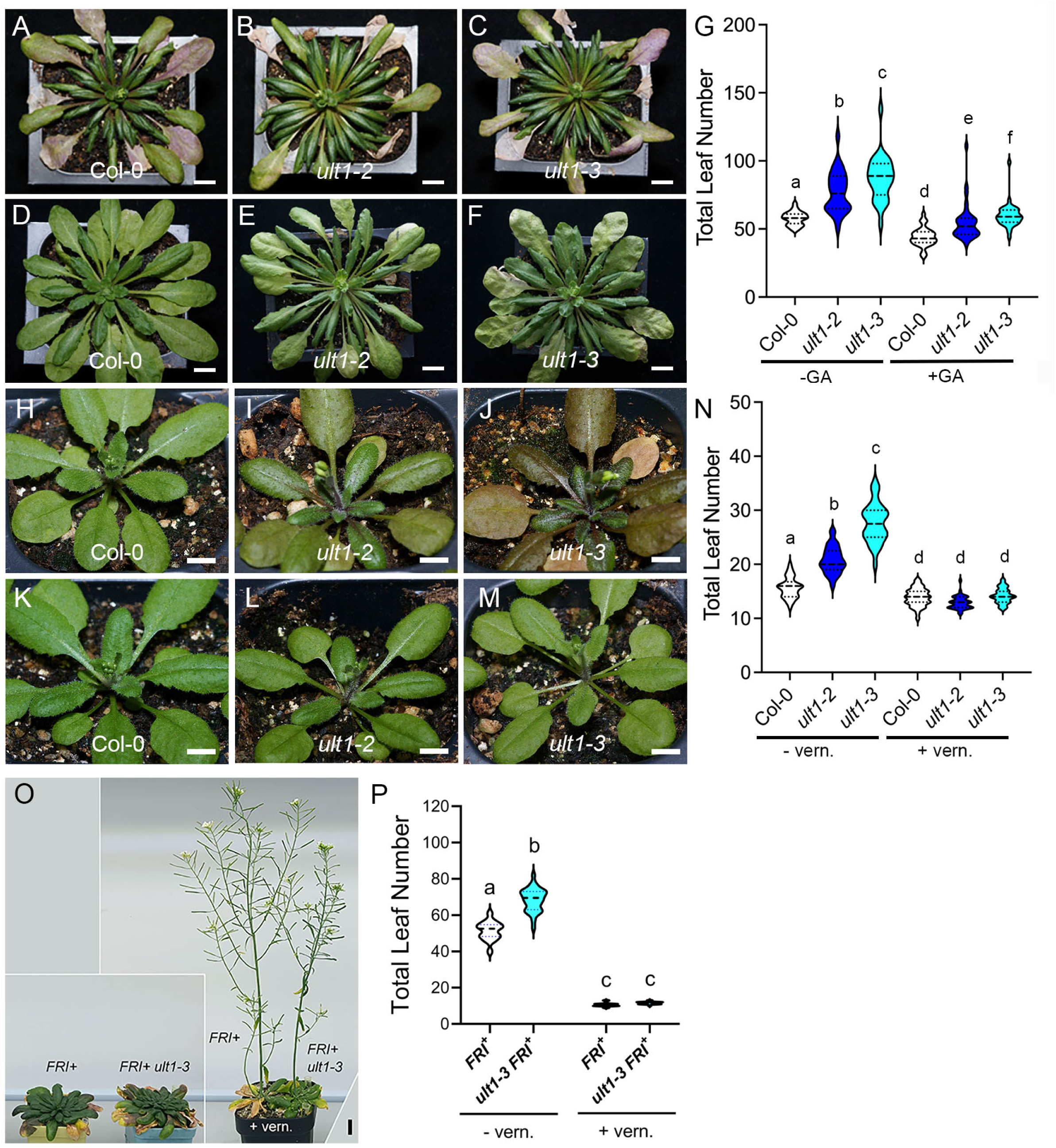
*ULT1* does not function in the gibberellic acid (GA) or vernalization pathways. **(A)** wild-type Col-0, **(B)** *ult1-2* and **(C)** *ult1-3* plants grown under SD conditions in the absence of GA. **(D)** wild-type Col-0, **(E)** *ult1-2* and **(F)** *ult1-3* plants grown under SD conditions in the presence of GA. **(G)** Mean total leaf number in the absence (-GA) or presence (+GA) of GA. **(H)** wild-type Col-0, **(I)** *ult1-2* and **(J)** *ult1-3* plants grown to bolting under LD conditions without cold treatment. **(K)** wild-type Col-0, **(L)** *ult1-2* and **(M)** *ult1-3* plants grown under LD conditions after 38 days of cold treatment. **(N)** Mean total leaf number in the absence (-vern.) or presence (+vern.) of cold treatment. **(O)** Left panel shows *FRI+* and *FRI+ ult1-3* plants grown to bolting under LD conditions without cold treatment and right panel shows *FRI+* and *FRI+ ult1-3* plants after 4 weeks (+vern.) of cold treatment. **(P)** Mean rosette leaf number of *FRI*+ and *FRI*+ *ult1-3* plants in the absence (-vern.) or presence (+vern.) of cold treatment. Lower case letters indicate statistically significant differences (P < 0.001). n ≥16. Scale bar, 1 cm.

Vernalization, or prolonged exposure to cold temperatures, accelerates the ability of Arabidopsis plants to flower as temperatures rise, due to gradual epigenetic repression of the *FLC* locus (Berry and Dean 2015). To determine if *ULT1* acts in the vernalization pathway we exposed wild-type and *ult1* seeds to 38 days of cold treatment at 4°C before growing them in LD conditions. The delay in *ult1-2* and *ult1-3* flowering under LD (Figure 2H-J and 2N) was completely corrected by the cold treatment (Figure 2K-N), indicating that *ult1* mutants respond normally to vernalization. Next we vernalized L*er* and *ult1-3* plants containing an active version of *FRIGIDA* (*FRI+*) that encodes a main activator of *FLC* in natural Arabidopsis accessions (Michaels and Amasino 1999). Like AP mutants, *FRI+* plants express high levels of *FLC* transcription that delays flowering under both LD and SD conditions but they show a strong response to vernalization (Lee and Amasino 1995; Michaels and Amasino 1999). To explore whether vernalization can promote earlier flowering in *ult1* mutants carrying a functional *FRI+* allele, we analyzed the effect of no cold treatment or 4 weeks cold treatment on the flowering time of L*er FRI+* and *ult1-3 FRI+* plants. We determined that in the absence of cold, *ult1-3 FRI+* plants flowered significantly later than L*er FRI+* plants, whereas after 4 weeks cold treatment the two genotypes flowered at the same time (Figure 2O and 2P). Therefore, *ult1* plants respond normally to vernalization even in the presence of high *FLC* levels conferred by the *FRI+* allele. Collectively our results showing that *ULT1* promotes flowering independently of photoperiod, temperature and hormone signaling are consistent with a function for *ULT1* in the autonomous flowering pathway.

To determine the molecular basis for the *ULT1* flowering time phenotype, we analyzed the expression of key regulatory determinants of the Arabidopsis floral transition by performing reverse transcription-quantitative polymerase chain reaction (RT-qPCR) experiments on wild-type L*er* and *ult1* plants grown for 14 days under LD conditions. At this 14 DAG time point L*er* but not *ult1-3* plants have undergone the floral transition. The main target of the photoperiod pathway is the *CONSTANS (CO)* transcription factor gene (Putterill et al. 1995). In agreement with our observation that *ULT1* promotes flowering independently of day length, we found that *CO* mRNA levels were unaltered in *ult1-3* plants (Figure 3A). In contrast, the expression of *FLC*, the floral repressor gene that is the key target of the autonomous pathway (Wu et al. 2020), was significantly up-regulated in *ult1-3* plants (Figure 3B), consistent with the *ult1* late flowering phenotype. Two *FLC* paralogues, *MAF4* and *MAF5*, were also up-regulated in *ult1-3* plants (Figure 3C and 3D); however, expression of the FLC complex component SVP was unaltered (Figure 3E). These data show that ULT1 is required to down-regulate the expression of multiple floral repressor genes during vegetative development.

**Figure 3.**
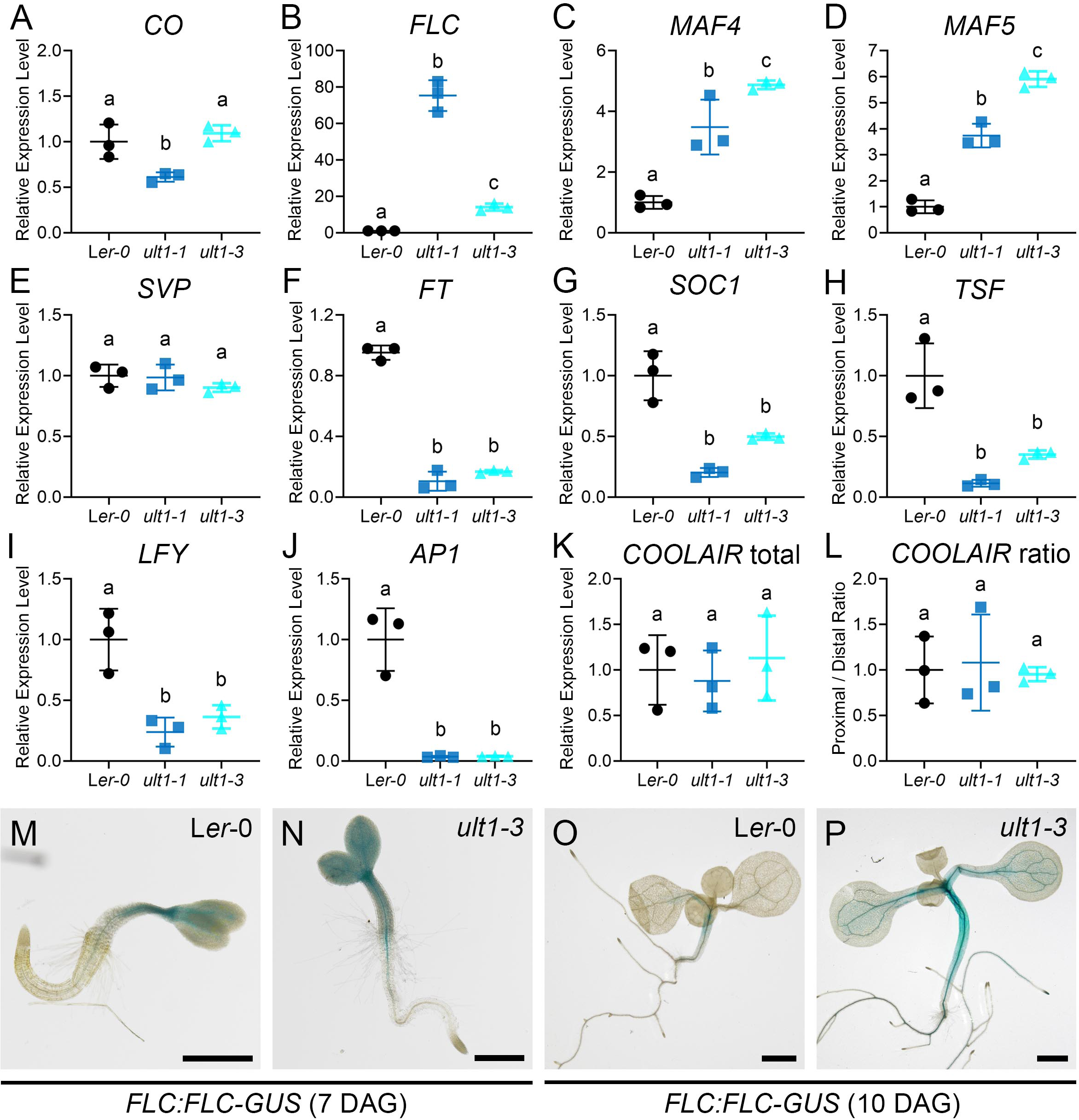
ULT1 regulates the expression levels of key flowering time genes. **(A-L)** Expression levels of **(A)** *CO*, **(B)** *FLC,* **(C)** *MAF4,* **(D)** *MAF5*, **(E)** *SVP,* **(F)** *FT*, **(G)** *SOC1*, **(H)** *TSF*, **(I)** *LFY*, and (J) *AP1*, genes as well as **(K)** total *COOLAIR* transcripts and **(L)** the ratio of proximal to distal *COOLAIR* transcripts relative to the *eIF4* reference gene in 14 DAG wildtype L*er, ult1-1* and *ult1-3* plants. Values indicate mean ± standard deviation for three biological replicates. Lower case letters indicate statistically significant differences (P < 0.05). **(M-P)** *FLC:FLC*-*GUS* reporter activity in a **(M)** 7 DAG L*er*, **(N)** 7 DAG *ult1-3*, **(O)** 10 DAG L*er* and **(P)** 10 DAG *ult1-3* seedling. Scale bar, 500 μm.

FLC functions as a direct repressor of the floral integrator genes *FT* and *SOC1* (Michaels et al. 2005; Helliwell et al. 2006), which induce flowering in response to various cues (He et al. 2020). Consistent with the *ult1* late flowering phenotype, the expression levels of both *FT* and *SOC1* were significantly reduced in *ult1-3* plants (Figure 3F and 3G), as was the expression of *TWIN SISTER OF FT (TSF)* (Figure 3H), which encodes as a floral integrator that acts partially redundantly with FT (Yamaguchi et al. 2005; Jang et al. 2009). Finally, two key floral meristem identity genes, *LEAFY (LFY)* and *APETALA1 (AP1),* were significantly down-regulated (Figure 3I-J). In sum, ULT1 promotes the expression of various floral integrator gene and floral meristem identity genes to accelerate flowering in wild-type seedlings.

We next assessed whether ULT1 was sufficient to regulate the expression of key downstream target genes in the flowering time pathway. To address this question, we performed RT-qPCR on wild-type L*er*, *35S*:*ULT1* and *ult1-3* plants grown for 12 days under LD conditions. Consistent with the data we collected at 14 DAG, we found that *CO* mRNA levels were unaltered in *ult1-3* plants (Supplemental Figure S3). Furthermore, *ult1-3* plants displayed significant up-regulation of *FLC* expression, accompanied by down-regulation of *FT* and *SOC1* expression (Supplemental Figure S3). Conversely, *FLC* transcript levels were significantly reduced, and *FT* and *SOC1* transcript levels significantly elevated, in *35S*:*ULT1* plants (Supplemental Figure 3). Thus the amount of *FLC, FT* and *SOC1* transcripts all respond to the level of *ULT1* activity, indicating that *ULT1* is sufficient to regulate the expression of these key determinants of the floral transition. The most parsimonious explanation for our results is that ULT1 represses *FLC* transcription, which in turn allows for induction of *FT* and *SOC1* expression to accelerate flowering.

We further analyzed the regulation of *FLC* expression by ULT1 during the vegetative phase by examining *FLC* expression in *ult1-3* seedlings using a *FLC:FLC*-*GUS* reporter line (Bastow et al. 2004). At 4 DAG, the *FLC:FLC-GUS* reporter was active throughout the above-ground tissues of L*er* seedlings and in the root vasculature (Figure 3M). The pattern was similar in *ult1-3* seedlings (Figure 3N), although the GUS activity in the cotyledons appeared qualitatively stronger. By 10 DAG, *FLC:FLC-GUS* reporter activity was restricted to the vasculature in L*er* seedlings (Figure 3O). In contrast, *ult1-3* seedlings displayed stronger *FLC:FLC-GUS* activity in the vasculature than L*er* seedlings and retained *FLC:FLC-GUS* activity in the hypocotyl and cotyledon tissues (Figure 3P). These findings indicate that ULT1 limits *FLC:FLC-GUS* activity in developing seedlings, particularly as they approach the transition to flowering.

### *FLC* is the Key Direct Biological Target of ULT1 Regulation

Because *FLC* is the predominant floral repressor gene in Arabidopsis and is the main target of the AP, we examined whether the delayed flowering phenotype of *ult1* plants was caused by the mis-regulation of *FLC* transcription. We tested the role of *FLC* by generating double mutants between *ult1-3* and *flc-3*, a null allele for the *FLC* locus (Michaels and Amasino 1999). When grown under LD conditions, *flc-3* plants flowered significantly earlier than either wild-type Col-0 (Figure 4A) or *ult1-3* (Figure 4B) plants, with an average of 13.74 total leaves per plant (Figure 4C and 4E). The *ult1-3 flc-3* double mutant plants flowered nearly as early as *flc-3* plants, with an average of 15.06 total leaves (Figure 4D and 4E). In contrast, although *maf5-3* mutants, which are homozygous for a T-DNA null allele (Shen et al. 2014), were slightly early flowering than Col-0 and *ult1-3* plants (Supplemental Figure 4A-C, E), the mean number of total leaves produced by *ult1-3 maf5-3* plants (Supplemental Figure 4D, E) was indistinguishable from that of *ult1-3* plants. Therefore, *MAF5* mis-regulation does not contribute to the *ult1* late flowering phenotype. These results indicate that up-regulation of *FLC* transcription levels conditions the *ult1* late flowering phenotype.

**Figure 4.**
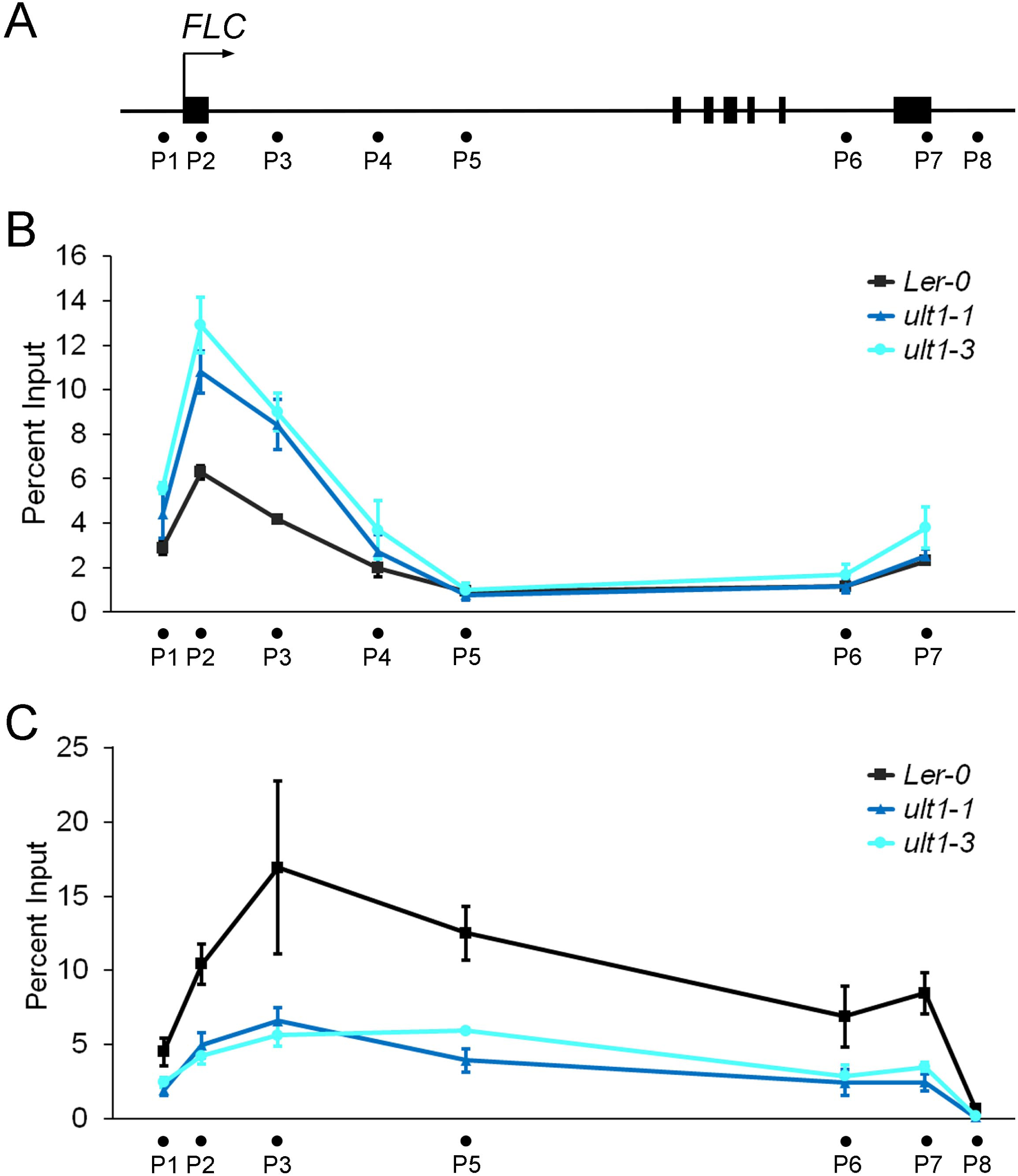
*FLC* is required to confer the *ult1* flowering time phenotype. **(A-D)** Top-down views of **(A)** wild-type Col-0, **(B)** *ult1-3,* **(C)** *flc-3* and **(D)** *ult1-3 flc-3* plants grown under LD conditions. **(E)** Mean total leaf number. Lower case letters indicate statistically significant differences (P < 0.001). n > 20. Scale bar, 1 cm.

Expression of the flowering integrator florigen gene *FT* is dramatically down-regulated in *ult1* seedlings, suggesting that ULT1 might promote flowering via regulation of *FT* transcription. To determine whether FT mis-regulation plays a role in conditioning the *ult1* late flowering phenotype we generated *ult1 ft* double mutants using the *ft-7* null allele in the L*er* accession (Onouchi et al. 2000). We observed that *ft-7* plants generated significantly more leaves than *ult1-3* or L*er* plants at flowering (Supplemental Figure 5), consistent with its primary role in promoting the floral transition. *ult1-3 ft-7* double mutant plants flowered much later than *ft-7* plants (Supplemental Figure 5), indicating that the two genes have a synergistic effect on flowering. We conclude that *ULT1* and *FT* function in separate genetic pathways to accelerate flowering in LD conditions. These data provide additional evidence for ULT1 regulating the floral transition independently of photoperiod. In sum, our results demonstrate that *FLC* and not *MAF5* or *FT* is the key biologically relevant target of ULT1 regulation during the floral transition.

Next we analyzed the molecular mechanism through which ULT1 regulates *FLC* transcription to accelerate flowering. ULT1 is a transcriptional regulator that can activate multiple developmental loci (Carles and Fletcher 2009; Pu et al. 2013; Tyler et al. 2019) and previously we have proposed that the protein, which carries a DNA-binding SAND domain, can act as direct regulator of gene expression by recruiting co-activator and RNA Pol II complexes for transcription initiation and elongation (Carles and Fletcher 2010). Because ULT1 functions in the AP, a straightforward mechanism through which ULT1 might regulate *FLC* expression is by activating other components of the AP that function as upstream repressors of *FLC* transcription. We tested this hypothesis by measuring AP gene expression levels in wild-type Col and *ult1-3* 12 DAG seedlings using RT-qPCR. We observed only a slight decrease in the mRNA expression levels of two AP pathway genes, *FLK* and *REF6* (Supplemental Figure 6). *REF6* was most affected, showing a 30% reduction in transcript levels in *ult1-3* compared to wild-type plants.

To determine if *ULT1* and *REF6* act in the same genetic pathway to regulate the floral transition, we generated double mutants between *ult1-3* and a null allele of *REF6, ref6-3*, and analyzed their flowering phenotype under LD conditions. We found that *ref6-3* plants (Supplemental Figure 7C) flower after producing significantly more leaves than either Col (Supplemental Figure 7A) or *ult1-3* plants (Supplemental Figure 7B). *ult1-3 ref6-3* plants (Supplemental Figure 7D) showed a synergistic increase in total leaf number compared to either single mutant (Supplemental Figure 7E), indicating that *ULT1* and *REF6* do not function in a single linear pathway to regulate flowering time. Coupled with the fact that *ult1-3* plants continue to transcribe all the AP genes to at least 70% of wild-type levels, these results disfavor a scenario in which *FLC* repression by ULT1 occurs indirectly through the induction of AP gene expression.

Multiple components of the AP promote flowering by activating expression of the *FLC* antisense transcript *COOLAIR* (Supplemental Figure 8A) (Liu et al. 2007; Swiezewski et al. 2009). The FCA, FY and FPA proteins are components of a *COOLAIR* RNA-processing machinery that promotes the usage of the proximal poly(A) site in *COOLAIR* (Liu et al. 2010). This *COOLAIR* transcript processing activity is associated with epigenetic silencing mechanisms that enrich H3K27me3 deposition at the *FLC* locus to repress its transcription (Swiezewski et al. 2009; Csorba et al. 2014; Tian et al. 2019). Therefore, we tested the hypothesis that ULT1 plays a role in regulating *COOLAIR* transcript abundance by analyzing total *COOLAIR* mRNA levels as well as those of the class I and class II *COOLAIR* splicing variants. We found no differences in total *COOLAIR* transcript levels nor changes in the ratio of the *COOLAIR* splicing variants between L*er* and *ult1-1* or *ult1-3* plants (Figure 3K-L). We also examined class I and class II *COOLAIR* transcript levels in 12 day old Col seedlings. Again, we detected no significant differences in *COOLAIR* transcript abundance between Col and *ult1-2* or *ult1-3* plants (Supplemental Figure 8). These findings are inconsistent with the hypothesis that ULT1 is involved in *COOLAIR* transcriptional control and instead suggest that ULT1 regulation of *FLC* transcription may occur via histone modification.

### ULT1 Regulates *FLC* Histone Methylation via Direct Binding to the *FLC* Locus

Previous studies showed that histone modifications such as H3K4me3 and H3K27me3 are crucial for the regulation of *FLC* transcription during vegetative development in response to the AP as well as to vernalization (Hepworth and Dean 2015; Wu et al. 2020). Considering that ULT1 can act together with trxG factors and PcG factors that modify histones, we examined the histone methylation patterns at the *FLC* locus in 21 DAG wild-type L*er* and *ult1-3* seedlings grown under LD conditions. We first analyzed the distribution of permissive H3K4me3 histone marks across the *FLC* locus. As expected, wild-type seedlings showed a peak of H3K4me3 enrichment around the translation start site (Figure 5B). This enrichment increased in *ult1-1* and *ult1-3* plants (Figure 5B and Supplemental Figure 9), revealing that ULT1 significantly decreases H3K4me3 enrichment at the *FLC* locus. H3K4me3 levels at the *FT, ACT2, ACT7* and *UBC* loci were unaltered in *ult1-1* plants (Supplemental Figure 9), indicating that ULT1 does not affect H3K4 trimethylation genome-wide. In contrast, H3K27me3 repressive mark levels were high across the *FLC* genomic region in 21 DAG L*er* plants, but were significantly reduced in *ult1-1* and *ult1-3* plants at almost all locations across the *FLC* locus (Figure 5C), particularly in the *FLC* promoter and first intron which are required for proper *FLC* regulation (Bastow et al. 2004; Kim et al. 2005). H3K27me3 levels at the *FT* locus, on the other hand, were unaltered in *ult1-1* and *ult1-3* plants but were strongly reduced in *emf2-10 vrn2-1* PcG double mutant plants (Supplemental Figure 10). These data indicate that the effect of ULT1 on H3K27me3 as well as H3K4me3 levels during flowering is specific to *FLC*. Thus ULT1 both reduces the accumulation of H3K4me3 active marks and promotes the accumulation of H3K27me3 repressive marks at the *FLC* locus, providing a chromatin context for the stable repression of *FLC* transcription during the floral transition.

**Figure 5.**
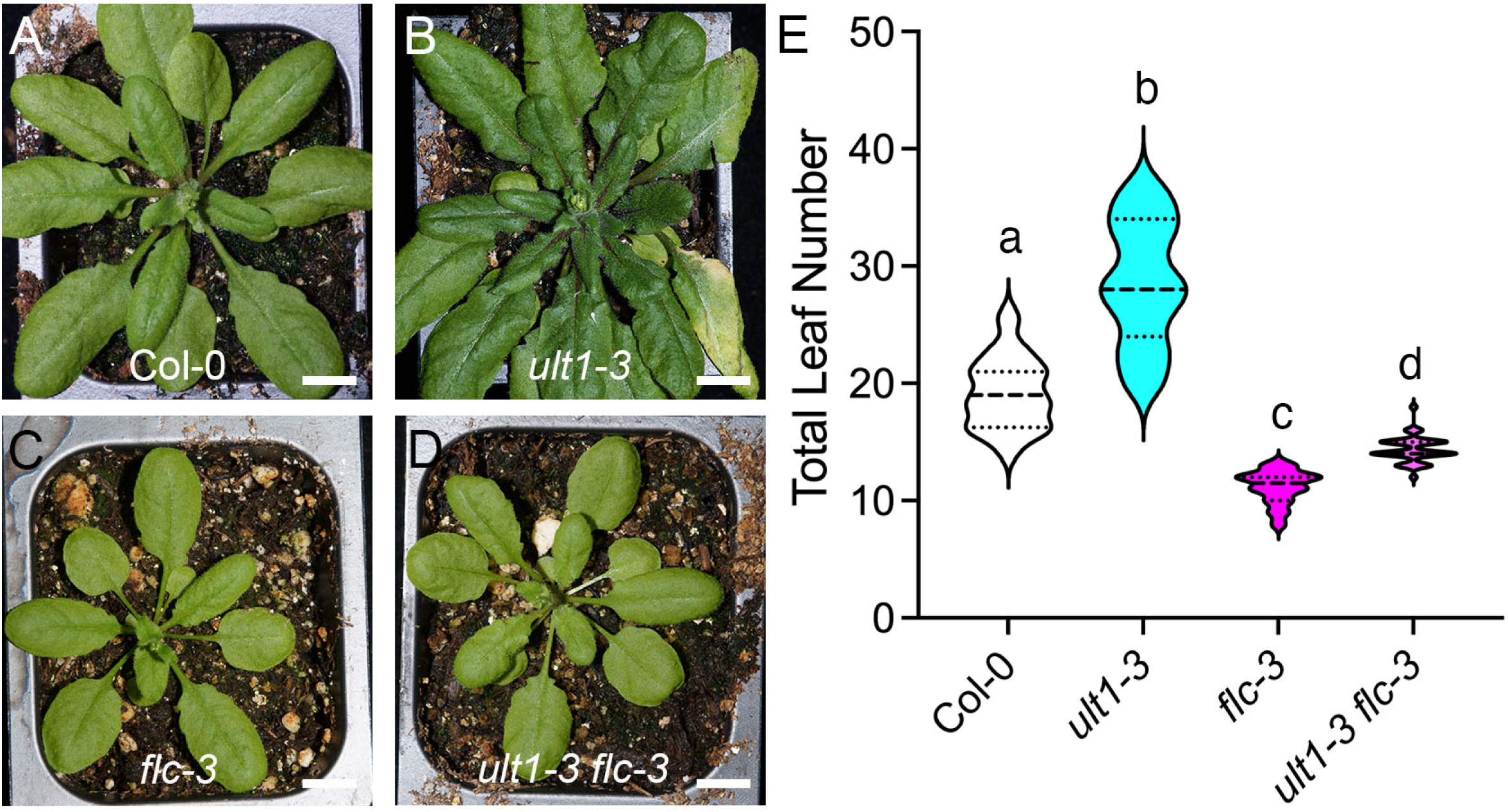
ULT1 regulates histone methylation levels at the *FLC* locus. **(A)** Schematic of the *FLC* genomic region with the large boxes indicating exons. Arrow indicates the *FLC* transcription start site (TSS). Black circles (P1-P8) indicate the positions of the primers used in the ChIP-qPCR analysis. **(B)** H3K4me3 levels across the *FLC* locus in L*er, ult1-1* and *ult1-3* 21 DAG plants. **(C)** H3K27me3 levels across the *FLC* locus in L*er, ult1-1* and *ult1-3* 21 DAG plants. Values indicate mean ± standard error for three biological replicates.

Because ULT1 can directly regulate gene transcription and histone modifications at target loci (Xu et al. 2018; Tyler et al. 2019) and does not appear to induce the activity of *FLC* repressor loci, we tested the hypothesis that ULT1 binds directly to the *FLC* locus (Figure 6A) to control its transcription. Chromatin immunoprecipitation followed by quantitative polymerase chain reaction (ChIP-qPCR) using an ULT1-3xMYC tagged protein driven by the *ULT1* promoter showed significant enrichment of ULT1-3xMYC protein at the P1 and P2 sites at the proximal promoter and first exon of *FLC* (Figure 6B). Maximal ULT1-3xMYC enrichment was observed at the P4 site in the first half of the first intron, insertions or mutations in which are known to affect *FLC* expression (Gazzani et al. 2003; Nasim et al. 2025). These results confirm binding of ULT1 to the *FLC* locus and are consistent with ULT1 acting as a direct negative regulator of *FLC* transcription during the floral transition.

**Figure 6.**
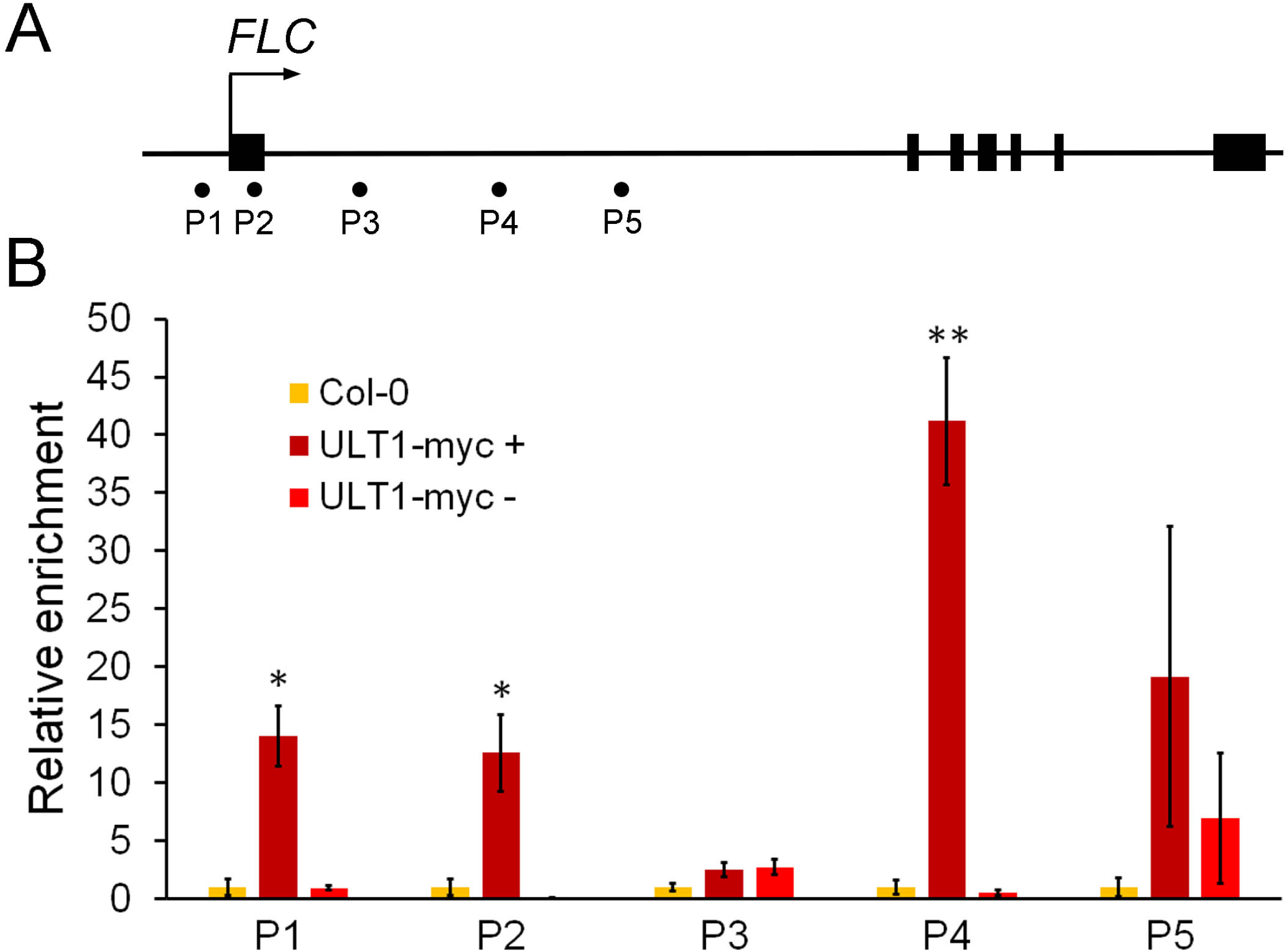
ULT1 directly binds to the *FLC* locus. **(A)** Schematic of the *FLC* genomic region. The large box indicates the first exon and the small grey box indicates the CME. Arrow indicates the *FLC* transcription start site (TSS). Black circles indicate the positions of the primers (P1-P5) used in the ChIP-qPCR analysis. **(B)** Relative enrichment of ULT1-3xMYC at five regions in and around the first exon of the *FLC* locus. Values indicate mean ± standard error for three biological replicates. * P < 0.05, ** P < 0.01.

### The PcG Methyltransferase Proteins SWN and CLF are Novel Direct ULT1 Interactors

Previous studies revealed that ULT1 functions in the nucleus as epigenetic co-regulator of the trxG protein ATX1 and the PcG accessory protein EMF1 (Carles and Fletcher 2009; Xu et al. 2018). Therefore, ULT1 may be directly involved in recruiting and/or stabilizing epigenetic regulatory complexes at the *FLC* locus during the floral transition. The known ULT1 interacting proteins ATX1 and EMF1 directly regulate *FLC* expression through the deposition of histone methylation marks, yet the loss of either *ATX1* or *EMF1* function causes earlier flowering (Aubert et al. 2001; Alvarez-Venegas et al. 2003) whereas *ult1* plants flower later than wild-type. That raises the possibility that the flowering repressive activity of ULT1 depends on physical interaction with additional unidentified epigenetic factors.

In order to identify novel ULT1 interaction partners, we used the full length ULT1 protein as bait in a yeast two hybrid (Y2H) screen of a commercial *Arabidopsis* cDNA library. Upon selection, we isolated a clone corresponding to locus At4G02020.1, which is annotated in the TAIR10 genome database as SWN, one of the three histone methyltransferase components of PRC2 (Chanvivattana et al. 2004). A targeted binary Y2H test using the full-length ULT1 and SWN proteins confirmed this interaction (Figure 7A). In subsequent pairwise Y2H tests we also detected an interaction between ULT1 and the SWN homologue CLF, and confirmed the interaction between ULT1 and EMF1 (Xu et al. 2018), although these interactions appeared to be weaker than that between ULT1 and SWN (Figure 7A). To test whether ULT1 can also directly interact with these proteins *in planta*, we performed firefly luciferase complementation imaging (LCI) assays in transiently transformed tobacco leaves (Figure 7B and 7C). We observed strong bioluminescence in the leaves when full-length ULT1 and SWN proteins were co-expressed as fusions with N- and C-terminal portions of luciferase (LUC), demonstrating that ULT1 and SWN proteins directly interact and reconstitute luciferase activity (Figure 7B). Co-expression of full-length CLF and ULT1 LUC fusion proteins also reconstituted luciferase activity (Figure 7C). Next, we assayed for an interaction between ULT1-nLUC and ULT1-cLUC proteins, based on the observation that the B-box motif mediates the multimerization of the OsULT1 protein (Roy et al., 2019), and determined that ULT1 can form homodimers (Figure 7C). Finally, we tested the interaction between CLF and FVE/MSI4, a DDB1 and CUL4-associated factor that is a key component of the AP and represses *FLC* transcription through its association with PRC2 (Pazhouhandeh et al. 2011). Co-expression of full-length CLF-nLUC and FVE-cLUC proteins likewise resulted in a physical interaction that produced strong luciferase activity (Figure 7B). These results indicate that the ULT1 proteins can not only homodimerize but also interact with both the SWN and CLF proteins, and that CLF can interact with FVE as well.

**Figure 7.**
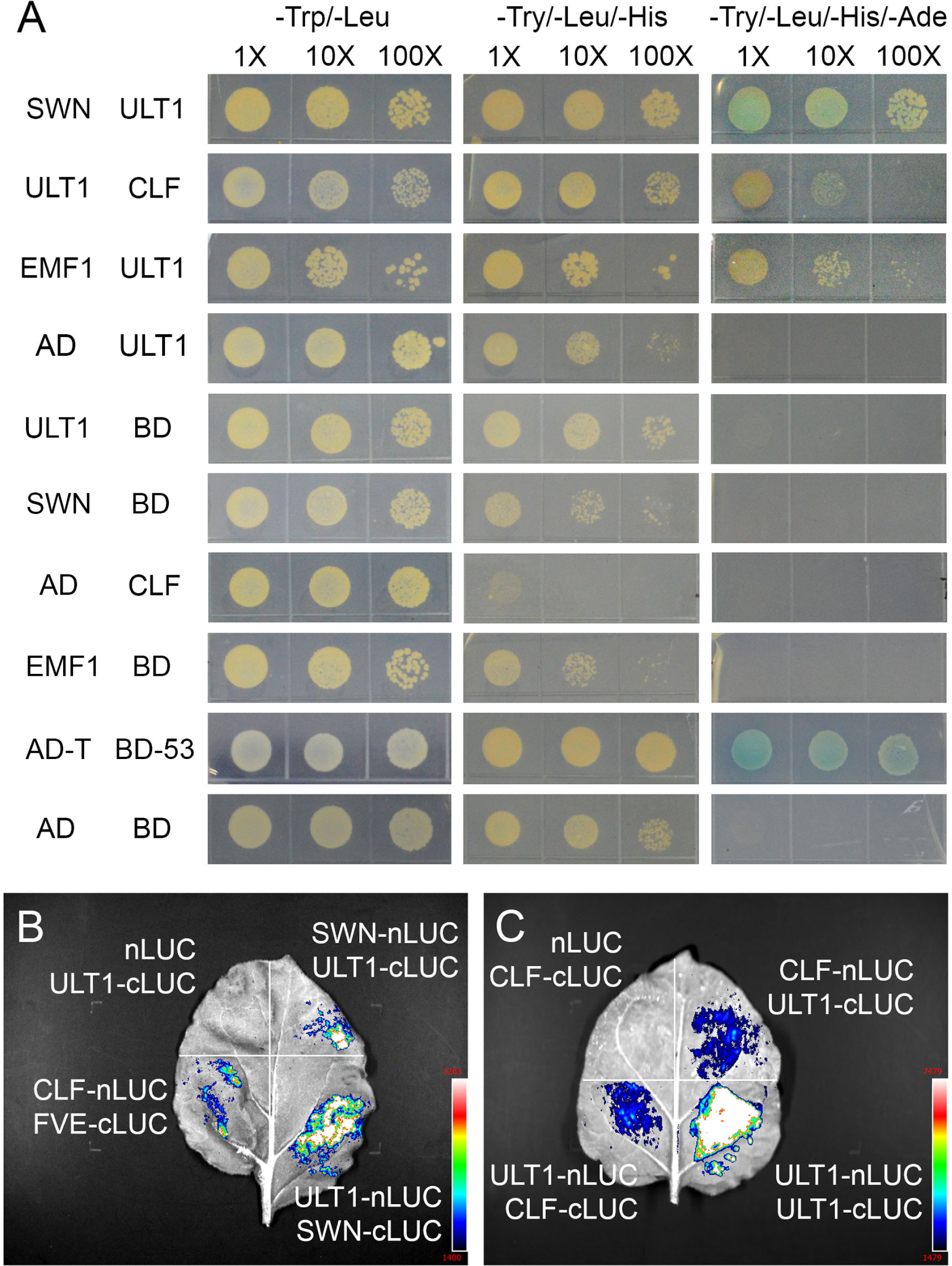
ULT1 physically associates with SWN and CLF proteins. **(A)** Yeast two hybrid assays testing for interaction between ULT1, SWN and CLF by monitoring the growth of yeast co-transformed with various prey and bait vectors on medium lacking Trp, Leu, His and Ade (-Try/-Leu/-His/-Ade) compared with control medium lacking only Trp and Leu (-Trp/-Leu), or Trp, Leu and His (-Try/-Leu/-His). AD is the activation domain empty vector and BD is the binding domain empty vector. **(B)** Luciferase complementation imaging (LCI) assays in *N. benthamiana* leaves co-expressing the negative control nLUC+ULT1-cLUC fusion protein combination, the positive control CLF-nLUC+FVE-cLUC fusion protein combination, the SWN-nLUC+ULT1-cLUC or the ULT1-nLUC+SWN-cLUC fusion protein combination in one of the four quadrants. **(C)** LCI assays in *N. benthamiana* leaves co-expressing the negative control nLUC+CLF-cLUC fusion protein combination or the ULT1-nLUC+CLF-cLUC, CLF-nLUC+ULT1-cLUC, or ULT1-nLUC+ULT1-cLUC fusion protein combination in one of the four quadrants.

We further investigated the interaction between ULT1 and the two PRC2 proteins by employing fluorescence resonance energy transfer fluorescence lifetime imaging microscopy (FRET-FLIM). We first assayed for interaction between SWN and ULT1 by transiently expressing SWN-mVenus in tobacco leaf epidermal cells (Figure 8A) and measuring its florescence lifetime (Figure 8G). Co-expression of SWN-mVenus with a negative control mCherry-NLS construct did not alter the mean florescence lifetime (Figure 8B and 8G). In contrast, co-expression of *SWN-mVenus* with *ULT1-mCherry* (Figure 8C) significantly reduced the mean florescence lifetime (Figure 8G), demonstrating a physical association between the two proteins. Next, we protoplasts either alone (Figure 8D) or co-expressed with *mCherry-NLS* (Figure 8E) or *ULT1-mCherry* (Figure 8F). Only the co-expression of the CLF and ULT1 fusion constructs resulted in a reduction in mean florescence lifetime (Figure 8H). These data demonstrate that ULT1 physically associates with the PRC2 components SWN and CLF *in planta*.

**Figure 8.**
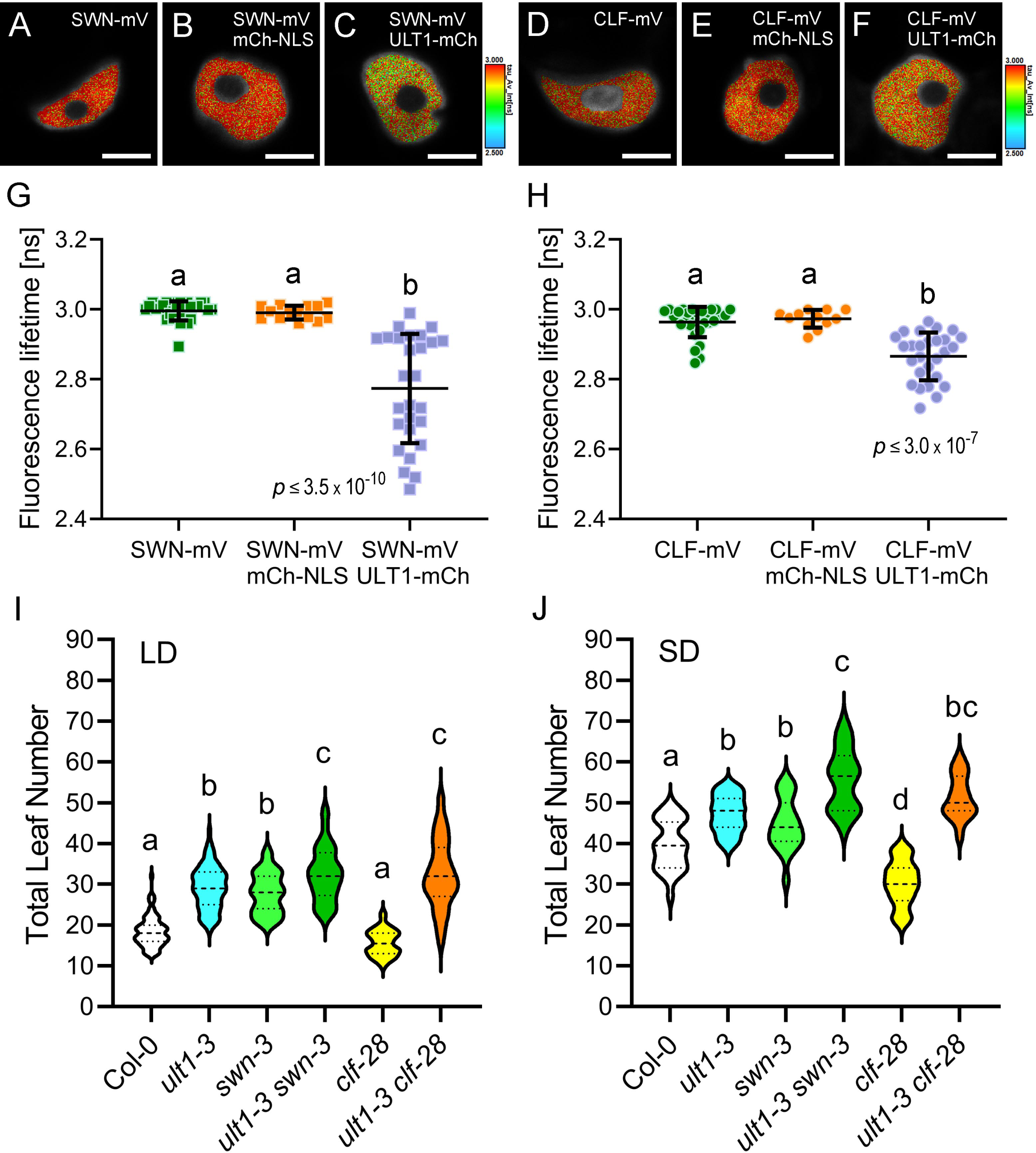
ULT1 physically associates with SWN and CLF and likely acts in the same pathway as SWN to promote flowering. Representative images of *N. benthamiana* nuclei showing the intensity-weighted fluorescence lifetime of (**A-C**) SWN-mVenus or (**D-F**) CLF-mVenus in the absence or presence of the indicated acceptor after fluorescence lifetime imaging microscopy measurements using a biexponential fitting model. The fluorescence lifetime in nanoseconds (ns) is color-coded with blue corresponding to a short lifetime and red corresponding to a long lifetime. Scale bar: 6 µm. (**G-H**) Quantification of the intensity-weighted fluorescence lifetime of (**G**) SWN-mV or (**H**) CLF-mV in the absence or presence of the indicated acceptor. n≥12. **(I)** Mean total leaf number of Col-0, *ult1-3, clf-28* and *swn-3* single and double mutants under LD conditions. n≥31. **(J)** Mean total leaf number of Col-0, *ult1-3, clf-28* and *swn-3* single and double mutants under SD conditions. n≥16. Lower case letters indicate statistically significant differences (P < 0.05).

Finally, we examined the biological relevance of the interaction between ULT1 and the CLF and SWN proteins by analyzing the flowering phenotypes of *ult1-3 clf-28* and *ult1-3 swn-3* double mutants grown in LD and SD conditions. As has been reported previously (Jiang et al. 2008; Lopez-Vernaza et al. 2012), *clf-28* null mutant plants flower one to two weeks earlier than wild-type Col plants in LD and SD conditions, although their rosette leaf number is equal to wild-type (LD; Figure 8I) or is only slightly reduced (SD; Figure 8J). This phenotype is due to *FT* misexpression through the loss of H3K27me3 repressive marks at the *FT* locus (Jiang et al. 2008; Lopez-Vernaza et al. 2012). In contrast, stronger de-repression of *FLC* in PcG double mutants prevents *FT* mis-expression (Müller-Xing et al. 2014). Under LD conditions, *ult1-3 clf-28* plants flower late after producing just a few more leaves than *ult1-3* plants (Figure 8I), indicating that *ult1* is largely epistatic to *clf* and that ULT1 activity is required for the *clf* early flowering phenotype. In contrast, *swn-3* null mutant plants flower later than wild-type under LD, producing the same number of leaves as *ult1-3* plants (Figure 8I). *ult1-3 swn-3* plants produce only a few more leaves than either *ult1-3* or *swn-3* plants (Figure 8I), a less-than-additive phenotype consistent with the two genes having predominantly overlapping functions in promoting flowering time. A similar outcome is observed in SD conditions, where *ult1* is epistatic to *clf* and *ult1-3 swn-3* plants have a less-than-additive flowering phenotype (Figure 8J). Taken together, our data indicate that ULT1 functions together with SWN- and CLF-containing PRC2 complexes to accelerate Arabidopsis flowering by directly regulating *FLC* transcription through the AP independently of photoperiod.

## DISCUSSION

Coordinating the transition to flowering in response to environmental and endogenous cues is essential for plant reproductive success. The regulation of the floral repressor gene *FLC* plays a key role in this process. In rapid-cycling Arabidopsis accessions such as Col and L*er*, *FLC* expression is high early during the vegetative phase and prevents premature activation of the floral inducers *SOC1* and *FT* and unseasonable flowering. As vegetative development progresses, environmental cues as well as the endogenous GA and autonomous pathways reduce *FLC* transcript levels via transcriptional and post-transcriptional mechanisms as well as epigenetic silencing (Kyung et al. 2022). This gradual reduction of *FLC* expression de-represses *SOC1* and *FT* transcription and induces flowering. Yet despite more than two decades of active study, new regulators of this crucial plant developmental transition continue to be discovered (Dai et al. 2024; Roque et al. 2024; Shao et al. 2025).

Here we have demonstrated that the transcriptional regulatory gene *ULT1* plays an important role in controlling the floral transition. We found that *ULT1* promotes the transition to flowering and is sufficient to induce flowering under LD and SD photoperiods (Figure 1, Figure S1-S2). In contrast, *ULT2* does not affect flowering time either alone or in the absence of *ULT1* (Figure 1), consistent with its lack of expression during the vegetative phase (Carles et al. 2005). *ult1* plants respond normally to both hormone treatment and vernalization (Figure 2), indicating that ULT1 is a novel component of the AP.

The main target of AP function is the *FLC* locus (Wu et al. 2020; Kyung et al. 2022). Consistent with this, our study demonstrates that *FLC* transcription is elevated in *ult1* plants (Figure 3 and Supplemental Figure 3) and that *flc* mutations fully suppress the *ult1* late flowering phenotype (Figure 4). In contrast, although transcription of the floral inducer gene *FT* is strongly down-regulated in *ult1* seedlings (Figure 3), *ft* and *ult1* mutations have a synergistic effect on flowering (Supplemental Figure 5). Thus the two genes function in separate genetic pathways and the reduction in *FT* expression in *ult1* seedlings is likely a secondary consequence of the up-regulation of its repressor, FLC. Taken together, these data indicate that *FLC* is the biologically relevant target of ULT1 regulation during the floral transition.

AP components control *FLC* expression via several potentially interconnected mechanisms. A large group of AP factors regulate the alternative processing of the antisense long noncoding RNA *COOLAIR*, which leads to recruitment of the histone demethylase FLD to *FLC* chromatin and the establishment of a repressive chromatin state (Wu et al. 2020; Kyung et al. 2022). The MSI-like WD40 protein FVE also contributes to epigenetic repression of *FLC* transcription by decreasing H3K4 trimethylation and H3 deacetylation at the *FLC* locus, likely by forming a complex with FLD and the histone deacetylase HDA6 (Yu et al. 2016). In addition, H3K27me3 repressive marks at *FLC* are strongly reduced in AP mutants, although it remains unclear how PRC2 proteins that catalyze H3K27 trimethylation are recruited to the *FLC* locus (Wu et al. 2020).

We conducted several sets of experiments to investigate the molecular mechanism through which ULT1 functions in the AP. First, we determined whether ULT1, which is known to activate developmental gene transcription, induces the expression of other AP genes that function as upstream repressors of *FLC* transcription. However, we found that ULT1 has little effect on AP gene transcription (Figure 5), and that the most significantly down-regulated gene, *REF6*, regulates the floral transition through a separate genetic pathway from ULT1 (Supplemental Figure 7). These data suggest that induction of AP gene expression is not the main mechanism through which ULT1 promotes flowering. Next, we analyzed *COOLAIR* transcript levels in *ult1* mutant plants. Neither class I nor class II *COOLAIR* transcript levels were significantly altered in the absence of ULT1 (Figure 4 and Supplemental Figure 8), disfavoring a mechanism in which ULT1 plays a role in the alternative processing of *COOLAIR* RNAs. Instead ULT1 reduces the accumulation of H3K4me3 active marks and promotes the accumulation of H3K27me3 repressive marks at the *FLC* locus, indicating that ULT1 regulates the floral transition via the epigenetic modulation of histone modifications as it does in other developmental contexts (Carles and Fletcher 2009; Pu et al. 2013; Xu et al. 2018; Xu et al. 2024). Interestingly, the effect of ULT1 on chromatin marks during the floral transition appears to be restricted to the *FLC* locus, because H3K4me3 and H3K27me3 levels at *FT* and other flowering time loci are unaltered in *ult1* plants (Supplemental Figure 9 and 10).

Our ChIP-qPCR results show that ULT1 protein binds sites in the proximal promoter, first exon and first intron of the *FLC* locus (Figure 6). Therefore, ULT1 can directly regulate *FLC* transcription. Yet because ULT1 lacks activation or repression domains (Carles et al. 2005) the protein must mediate its effects via physical association with other proteins. Using a Y2H screen to detect novel ULT1 interaction partners, we isolated a clone corresponding to the histone methyltransferase SWN and verified direct ULT1-SWN and ULT1-CLF interaction in further Y2H assays (Figure 7A). Further analysis using LCI assays and FRET-FLIM confirmed that ULT1 physically associates with both SWN and CLF *in planta* (Figure 7B-C and 8A-H). Genetic epistasis tests demonstrated that ULT1 and CLF function in the same genetic pathway, and that ULT1 and SWN pathways also largely overlap (Figure 8I-J). These results indicate that during vegetative development ULT1 plays a role in establishing and/or maintaining H3K27 trimethylation at the *FLC* locus through the recruitment of PRC2.

Our study identifies a role for the ULT1 transcriptional regulator in another key developmental transition. In addition to its function in regulating the germination switch from embryo to seedling fate (Xu et al. 2018), the floral determinacy switch from meristem to carpel fate (Carles et al. 2004; Carles and Fletcher 2009), and the shoot to root fate switch during *de novo* root regeneration (Tian et al. 2022), we show here that ULT1 also promotes the floral transition switch from vegetative to reproductive fate. In most of these contexts ULT1 has been shown to work in combination with distinct subsets of histone modifying factors: ATX1 and EMF1 in germination, ATX1 in floral determinacy, and CLF and SWN in the floral transition. The ability of ULT1 to physically associate with both trithorax group (ATX1) and Polycomb group (EMF1, CLF, SWN) factors, to increase or decrease gene expression, and to regulate the deposition of H3K4me3 and H3K27me3 marks has led to the proposal that ULT1 functions as a molecular epigenetic switch that regulates transcription activation and repression in a context-dependent fashion (Ornelas-Ayala et al. 2021). Such molecular epigenetic switches permit dynamic gene regulation in response to developmental or environmental cues, leading to rapid changes in cell states, and regulate developmental cell fate decisions in multicellular organisms (Gomez-Schiavon and Buchler 2019). Consistent with this notion, the human SAND domain-containing protein AIRE was recently shown to preferentially target genes with promoters in a poised state (Fang et al. 2024), permitting the rapid induction of transcription cascades that drive the elimination of self-reactive T cells from the immune system. The selective deployment of ULT1 as a component of an epigenetic switch consisting of stage- and tissue-specific DNA-binding transcription factors (Pires et al. 2014; Moreau et al. 2016) along with select chromatin modifying factors may thus play a key role in mediating plant cell fate transitions, including the floral induction process.

Finally, our study uncovers a novel molecular mechanism for PRC2 recruitment to the *FLC* locus during vegetative development in preparation for flowering. ULT1 proteins bind to the promoter and first intron of *FLC* and interact with the SWN/CLF histone methyltransferases to recruit PRC2 to the locus. There the PRC2 complex deposits H3K27me3 marks and associates via CLF with FVE, component of a histone deacetylation complex, coordinating these epigenetic modifications to establish a repressive chromatin environment and down-regulate *FLC* expression. Reduction in FLC levels then releases the floral integrators *SOC1* and *FT* from FLC-mediated repression and allows them to induce the floral transition. Further investigation will be required to determine whether ULT1-associated changes in H3K4me3 marks at the *FLC* locus also contribute to *FLC* down-regulation, and to identify the histone methyltransferases involved.

## MATERIALS AND METHODS

### Plant materials, growth conditions and flowering time analyses

All Arabidopsis mutants used in this study are in the Columbia (Col-0) or the Landsberg *erecta* (L*er-*0) background. The *ult1-1, ult1-2, ult1-3*, *ult2-2* and *ult2-3* alleles have been previously described (Fletcher 2001; Carles et al. 2005; Monfared and Fletcher 2014). The *ult1-3* allele was originally isolated in Col-0 and was backcrossed four times to L*er* prior to analysis in that background. The *ult2-4* allele is a T-DNA insertion mutation (SALK_206246C) in the first exon of *ULT2*, +180 base pairs downstream of the start codon, in the middle of the conserved SAND domain. The *35S:ULT1* (Carles and Fletcher 2009), *flc-3* (Michaels and Amasino 1999), *ft-7* (Onouchi et al. 2000), *ref6-3* (Noh et al. 2004), *clf-28* (Doyle and Amasino 2009), *swn-3* (Chanvivattana et al. 2004), and *emf2-10 vrn2-1* (Müller-Xing et al. 2014) lines have been previously described. Primer sequences for genotyping are listed in Supplemental Table 1.

Arabidopsis seeds were sown in soil consisting of 50% medium vermiculite and 50% Sunshine Mix #1, and stratified for 5 days at 4°C before being transferred to a growth chamber where they were grown under 90-100 μE m^-2^ s^-1^ of light at 21°C. For LD conditions a photoperiod of 16 hours light and 8 hours dark was used, and for SD conditions a photoperiod of 8 hours light and 16 hours dark was used. For the vernalization experiments, seeds were planted on moisturized soil and incubated in darkness at 4°C for either 28 or 38 days before being placed under LD conditions at 21°C. For the GA treatments, 100 μM GA_3_ in ethanol was sprayed on the plants once a week until flowering, with ethanol alone used as the untreated control, as described (Deng et al. 2007). At least 20 plants for each genotype were scored for the flowering time analyses, except for the L*er* and *35S:ULT1* SD experiment in which the L*er* n = 14.

### Plasmid construction

For the yeast two-hybrid constructs, the full-length *CLF*, *SWN*, *EMF1* and *ULT1* coding sequences (CDSs) were amplified using gene-specific primers (Supplemental Table 1). The *SWN*, *CLF* and *EMF1* PCR fragments were digested with *SfiI* and *XmaI*, and the *ULT1* PCR fragments were digested with *EcoRI* and *BamHI*. The resulting fragments were inserted into the *pGADT7* and/or *pGBK7* vectors, respectively.

For the luciferease complementation imaging assay constructs, the full-length *ULT1* CDS was amplified using gene-specific primers (Supplemental Table 1) and cloned into the *pCAMBIA1300-nLUC* and *pCAMBIA1300-cLUC* vectors through homologous recombination using a one-step cloning kit (Vazyme, C112). Similarly, the full-length *CLF*, *SWN* and *FVE*CDSs were amplified using gene-specific primers (Supplemental Table 1) and cloned into the *pCAMBIA1300-nLUC* and *pCAMBIA1300-cLUC* vectors. All clones were screened and verified by restriction endonuclease test digestions and sequencing.

For the FRET-FLIM constructs, the full-length *ULT1* CDS was amplified using gene-specific primers (Supplemental Table 1) and cloned into the empty *pGGC000* entry vector at the *BsaI* site in a GreenGate reaction (Lampropoulos et al. 2013). After a second GreenGate reaction using *pGGZ001* as the destination vector, we obtained the *35S:ULT1-N-Venus* and XVE:*ULT1-mCherry* constructs. Similarly, the full-length *CLF* and *SWN* CDS were amplified using gene-specific primers (Supplemental Table 1) and cloned into the empty *pGGC000* entry vector, which after a second GreenGate reaction was used to obtain the *XVE:CLF-mVenus*, *XVE:SWN-mVenus*, *35S:CLF-C-Venus* and *35S:SWN-C-Venus* constructs. The *XVE:mCherry-NLS* construct has been described (Strotmann et al. 2025). All clones were screened and verified by restriction endonuclease test digestions and sequencing.

### Quantitative RT-PCR

Total RNA from three biological replicates was extracted using TRIZOL (Invitrogen) and cDNA was synthesized using RevertAid reverse transcriptase (Thermo Fisher). Real-time RT-qPCR was performed using SYBR Green I in a LightCycler 480 (Roche) thermocycler. Primer sequences are listed in Supplemental Table 1.

### Chromatin immunoprecipitation-quantitative PCR

ChIP assays were performed as described previously (Müller-Xing et al. 2014). The chromatin was fragmented to an average length of 200-400bp by sonication. Anti-tri-methylated histone H3K27 antibody (Abcam; ab6002) and anti-tri-methylated histone H3K4 antibody (Abcam; ab8580) were used for immunoprecipitation. DNA was recovered by Phenol:chloroform:Isoamyl Alcohol (25:24:1) and analyzed by ChIP-qPCR. Primer sequences are listed in Supplemental Table 1.

### Yeast two-hybrid screens and assays

Yeast two-hybrid (Y2H) assays were performed according to the Yeast Protocols Handbook and the Yeastmaker^TM^ Yeast Transformation System 2 (Cat. Nos. 630439, 630489 and 630490, Clontech^®^). For large-scale Y2H screening, a normalized library was used (Mate & Plate^TM^ Library-Universal *Arabidopsis*, Cat. No. 630487, Clontech^®^). The library was transformed into yeast strain Y187 (MATα), which can be readily mated to a MATa GAL4 reporter strain such as AH109 or Y2H Gold. The Matchmaker Gold Yeast Two-Hybrid System (Clontech^®)^ was used to screen the library. In brief, the *ULT1* cDNA bait was cloned into the pGBKT7 vector (GAL4 DNA-BD) and transformed into the Y2H Gold yeast strain. The mixture of the Y2H Gold bait strain and prey strain Y187 was cultured at 30℃ for 105 min. Vacuum filtration was used to collect the zygotes on YPDA medium incubated at room temperature overnight. Under a light microscope typically three-lobed structures can be seen clearly and spread on stringent SD (Synthetically Defined) selection medium (–Leu/–Trp/–His/3AT). Single colonies were picked and then grown on selective SD medium (–Leu/–Trp/–Ade/–His/X-α-gal/3AT). DNA from blue yeast colonies that survived this stringent selection was extracted and sequenced using the Matchmaker™ AD LD-Insert Screening Amplimer Set Primer (Cat. No. 630433, Clontech^®^). To verify genuine positive interactions, prey plasmids (pGADT7 DNA-AD containing the selected genes) were co-transferred with the corresponding bait plasmids into the Y2H system (Y2H Gold strain) on SD medium (-Leu/-Trp/-Ade/-His/X-α-gal).

For the binary Y2H interaction assays, the direct interaction of two proteins was investigated by co-transformation of the respective plasmids into the yeast strain AH109 or Y2H Gold (Clontech^®^), followed by selection of transformants on medium lacking Leu and Trp at 30℃ for 3 days. The selected strains were then dotted on −LT, −LTH or −LTAH medium (lacking Leu, Trp, and/or His and Ade) for growth selection and lacZ activity testing of interacting clones.

### Transient transformation of *Nicotiana benthamiana*

For transient expression in *Nicotiana benthamiana* (*N. benthamiana*), the *Agrobacteria* strain GV3101::pMP50 was used. The bacteria were grown overnight in 5 ml dYT medium containing the appropriate antibiotics at 28°C with shaking. First, the bacteria were centrifuged for 10 min at 4,000 rpm and 4 °C before the pellet was resuspended in infiltration medium (5% sucrose (w/v), 0.01% MgSO_4_ (w/v), 0.01% glucose (w/v) and 450 µM acetosyringone) to an optical density OD_600_ of 0.4. To ensure sufficient expressing cells, *Agrobacteria* were mixed with an *Agrobacterium* strain carrying the p19 silencing repressor and a second *Agrobacterium* strain carrying a different construct for co-expression of ULT1-mCherry. Subsequently, the cultures were incubated for 1 h at 4°C. *N. benthamiana* plants were sprayed with water and kept under high humidity prior to infiltration, to trigger stomatal opening and thereby allow effortless infiltration. The abaxial side of the leaf was infiltrated using a syringe without a needle. Expression was induced 2-4 days after infiltration by spraying a 20 µM β-estradiol solution containing 0.1% Tween®-20 (v/v) to the abaxial side of the leaf. Depending on the expression level, FLIM measurements were performed 10-16 h after induction.

### FRET-FLIM measurements

FRET-FLIM was performed in transiently expressing epidermal leaf cells of 3 to 4 weeks old *N. benthamiana* plants using an inverted LSM780 (Carl ZEISS GmbH) equipped with additional mVenus was chosen as donor and excited at 485 nm with a pulsed laser diode and ∼ 1 µW laser power at the objective (40 x C-Apochromat/1.2 Corr W27, ZEISS) and a frequency of 32 MHz, and detected using two τ-SPAD single photon counting detectors in perpendicular and parallel orientation. Photons were collected over 40 frames at 256x256 pixels per frame, a pixel dwell time of 12.6 µs and a digital zoom of 8. Prior to image acquisition, a calibration routine was performed. To test system functionality, fluorescence correlation spectroscopy (FCS) measurements of deionized water and Rhodamine110 were acquired. Additionally, monitoring the decay of erythrosine B in saturated potassium iodide served as instrument response function (IRF) to correct the fitting for system specific time shift between laser pulse and data acquisition. Analysis of FRET-FLIM measurements was performed in SymPhoTime64 (PicoQuant GmbH) using the ‘Grouped FLIM’ analysis tool. For donor-only samples, a monoexponential decaying model was used, whereas for samples that contain donor as well as acceptor a biexponential fit was chosen. Graphs were generated using the R and Prism 9 software packages.

### Imaging

Photographs were taken either with digital SLR cameras (Nikon D7200 or D3200, Japan), a dissecting microscope with a 5 Mega Pixel digital camera (Motic K-500L, China), or a Nikon SMZ25 stereomicroscope. Image processing was performed using Fiji ImageJ utilizing LUT ‘Magenta’. Digital photographs and graphics were collated with Microsoft PowerPoint or Adobe Photoshop and adjusted as described (Müller-Xing et al. 2014). Briefly, adjustment involved only image exposure using adjustment levels and sharpening using unsharp mask.

### Statistical analysis

All qPCR values represent the mean ± standard error. Statistical analysis of the flowering time experiments was performed using one-way ANOVA with *post-hoc* Tukey’s test for multiple comparisons. Statistical analysis of the *35S:ULT1* SD experiment was performed using Welch’s two-tailed t-tests. One-tailed Student’s *t* tests, resulting in *P* values, were employed to assess statistical significance between all other pairs of values. Statistical testing of FRET-FLIM data was performed using the R software package. Data were tested for homogeneity of variances (α = 0.05) and normal distribution (α = 0.05) using Levene’s test and Shapiro test, respectively. Because some data sets did not show homogeneity of variances or normal distribution, a non-parametric Kruskal-Wallis with post-hoc Dunn’s test (α = 0.05) with Benjamini and Hochberg correction was used.

## ACCESSION NUMBERS

CLF, At2g23380; EMF1, At5g11530; EMF2, At5g51230; FLC, At5g10140; FVE, At2g19520; FT, At1g65480; REF6, At3g48430; SWN, At4g02020; ULT1, At4g28190; ULT2, At2g20825; VRN2, At4g16845.

## ACKNOWLEDGEMENTS

We are grateful to Richard Amasino and Sara Farrona for discussions and supplying seeds, and all members of our groups for helpful discussions. We would like to also acknowledge the Center for Advanced Imaging (CAi) at Heinrich-Heine-University Düsseldorf for providing access to the Zeiss LSM780 microscope. This work was supported by the US National Science Foundation (IOS-105020 to JCF), the US Department of Agriculture (CRIS 2030-21000-048-00D to JCF), the National Natural Science Foundation of China (Project No. 32460150, 31640054 and 31771602 to RM-X), the Jiangxi Province Recruitment Program of Foreign Experts (Project No jxsq2023104003 to RM-X), and a HORIZON-MSCA Fellowship (101155273 to QX). Funding for instrumentation: Zeiss LSM780: DFG-INST 208/551-1 FUGG.

## AUTHOR CONTRIBUTIONS

Experiments were designed by RM-X, JCF, LP and ES and conducted by QX, ES, JT, YX, TY, VS, XL, YZ, JS, LP, JCF, RM-X, and XW. Data were analyzed by RM-X, JCF, QX, LP, ES and JT. YS provided important materials. The manuscript was written by JCF, RM-X and QX with the help of JT, and edited by YS. All authors read and approved the final manuscript.

