## Supplementary material for "The bivalent epigenetic regulator ULTRAPETALA1 promotes the Arabidopsis floral transition via recruitment of Polycomb histone methyltransferases for H3K27me3 deposition at the *FLC* locus": Supp data

Running title: ULT1 regulation of floral transition

[1] Jiangxi Provincial Key Laboratory of Plant Germplasm Innovation and Genetic Improvement, Lushan Botanical Garden, Chinese Academy of Sciences, Jiujiang 332900, China
[2] Plant Epigenetics and Development, Lushan Botanical Garden, Chinese Academy of Sciences, Nanchang, 330114, China
[3] College of Life Science, Nanchang University, Nanchang 330047, China
[4] Plant Gene Expression Center, United States Department of Agriculture - Agricultural Research Service, Albany, CA 94710, USA
[5] Department of Plant and Microbial Biology, University of California, Berkeley, Berkeley, CA 94720, USA
[6] Institute for Developmental Genetics, Heinrich-Heine University, Düsseldorf 40225, Germany
[7] Institute for Plant Developmental Genetics, Goethe-University, Frankfurt 60438, Germany
[8] Biotechnology Research Institute, Chinese Academy of Agricultural Sciences, Beijing 100081, China

[†] These authors contributed equally to this work

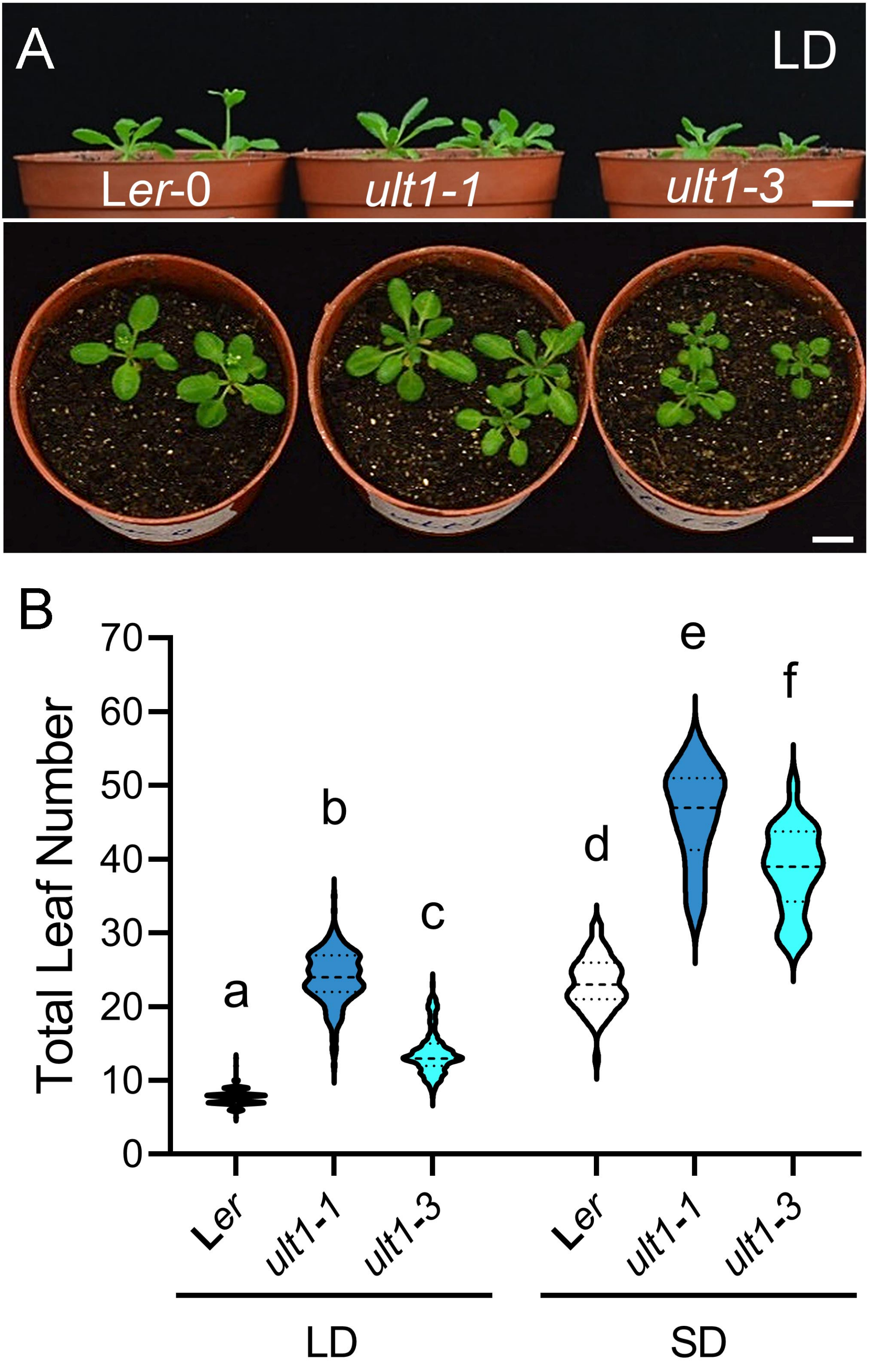
A
LD
Ler-0
ult1-1
ult1-3
B
Total Leaf Number
70
60
50
40
30
20
10
0
a
b
c
d
e
f
Ler
ult1-1
ult1-3
Ler
ult1-1
ult1-3
LD
SD

**Supplemental Figure 1.** *ULT1* alleles in the L*er* background are late flowering under long day (LD) and short day (SD) conditions. (**A**) Side views (upper panel) and top-down views (lower panel) of wild-type L*er-0, ult1-1* and *ult1-3* plants grown under LD conditions. (**B**) Mean total leaf number under LD and SD conditions. **** $P < 0.0001$. $n \geqslant 14$. Scale bar, 1 cm.

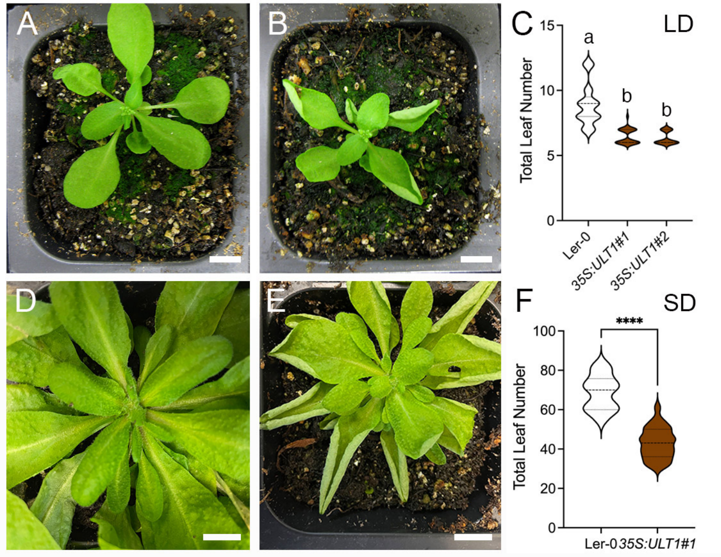
A
B
C
LD
15
10
5
0
Total Leaf Number
a
b
b
Ler-0
35S:ULT1#1
35S:ULT1#2
D
E
F
SD
****
100
80
60
40
20
0
Total Leaf Number
Ler-0
35S:ULT1#1

**Supplemental Figure 2.** ULT1 is sufficient to promote flowering independent of photoperiod. **(A-B)** Top-down views of **(A)** wild-type L*er-0* and **(B)** *35S:ULT1* plants grown under LD conditions. **(C)** Mean total leaf number under LD conditions. Data from two independent *35S:ULT1* lines are shown. **(D-E)** Top-down views of **(D)** L*er-0* and **(E)** *35S:ULT1* plants grown under SD conditions. **(F)** Mean total leaf number under SD conditions. Lower case letters indicate statistically significant differences ($P < 0.001$). **** $P < 0.001$. $n = 14$-27. Scale bar, 1 cm.

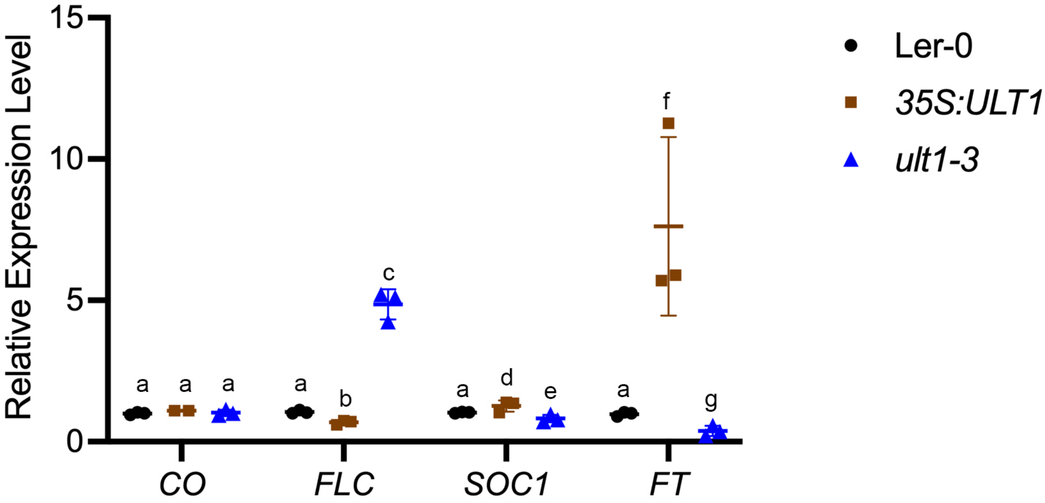

Relative Expression Level
15
10
5
0
a
a
a
a
b
c
a
d
e
a
f
g
CO
FLC
SOC1
FT
Ler-0
35S:ULT1
ult1-3

**Supplemental Figure 3.** *ULT1* is sufficient to regulate key flowering time integrator genes. mRNA expression levels of the flowering time regulatory genes *CO, FLC, FT* and *SOC1* in 12 DAG wild-type L*er-0*, *35S:ULT1* and *ult1-3* plants grown under LD conditions. Values represent mean ± standard error for three biological replicates. Lower case letters indicate statistically significant differences ($P < 0.001$).

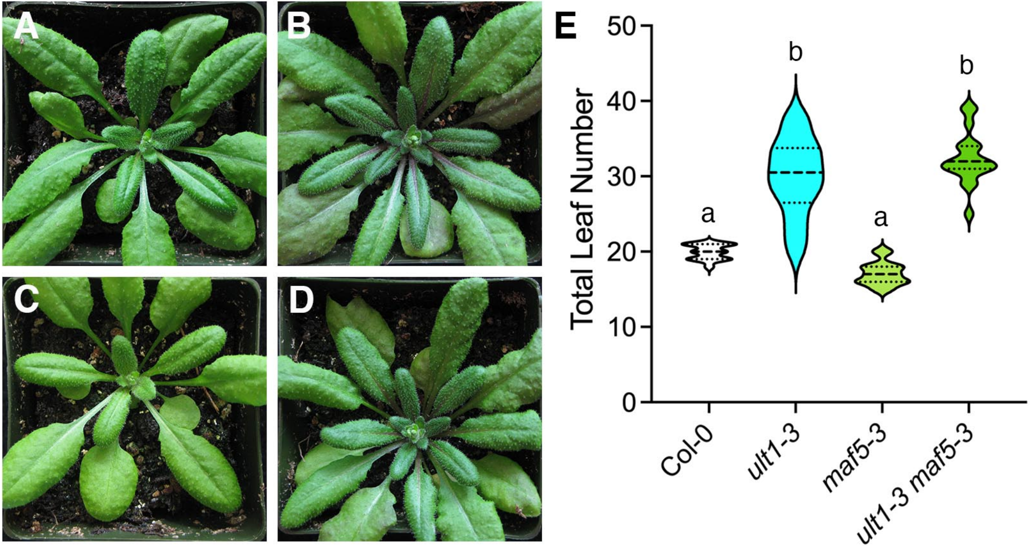
A
B
C
D
E
Total Leaf Number
50
40
30
20
10
0
a
b
a
b
Col-0
ult1-3
maf5-3
ult1-3 maf5-3

**Supplemental Figure 4.** *MAF5* mutations fail to rescue the *ult1* flowering time phenotype. **(A-D)** Top-down views of **(A)** wild-type Col-0, (**B**) *ult1-3,* **(C)** *maf5-3* and **(D)** *ult1-3 maf5-3* plants grown under LD conditions. **(E)** Mean total leaf number. Lower case letters indicate statistically significant differences ($P < 0.001$). $n > 20$. Scale bar, 1 cm.

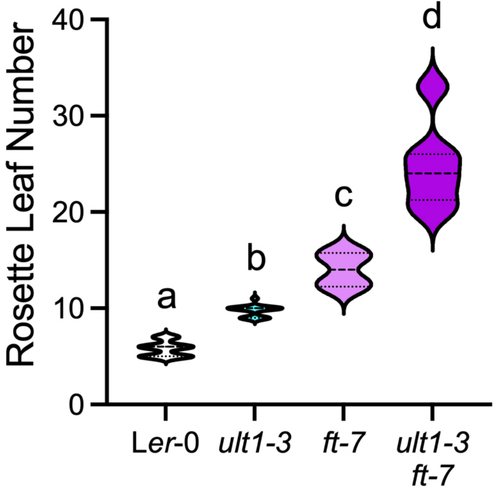

Rosette Leaf Number
40
30
20
10
0
a
b
c
d
Ler-0
ult1-3
ft-7
ult1-3
ft-7

**Supplemental Figure 5.** *FT* mutations enhance the *ult1* flowering time phenotype. Mean rosette leaf number of L*er-0, ult1-3, ft-7* and *ult1-3 ft-7* plants grown under LD conditions. Lower case letters indicate statistically significant differences ($P < 0.01$). n $\geqslant$44.

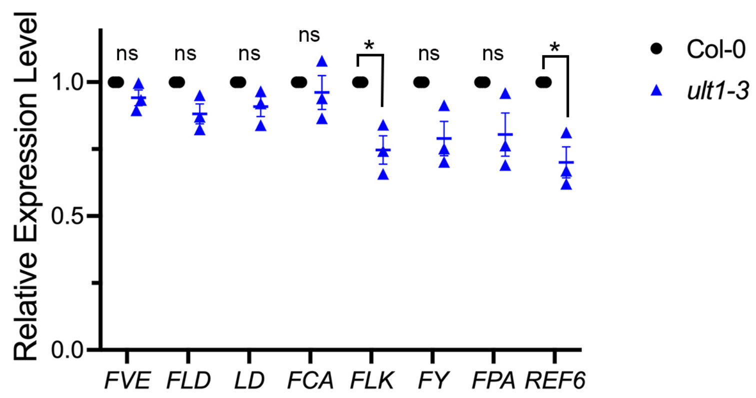

Relative Expression Level
0.0
0.5
1.0
ns
ns
ns
ns
*
ns
ns
*
FVE
FLD
LD
FCA
FLK
FY
FPA
REF6
Col-0
ult1-3

**Supplemental Figure 6.** ULT1 has little effect on autonomous pathway (AP) gene expression. mRNA expression levels of AP genes in 12 DAG wild-type Col-0 and *ult1-3* plants grown under LD conditions. Values represent mean ± standard error for three biological replicates. * $P < 0.05$. ns = not significant.

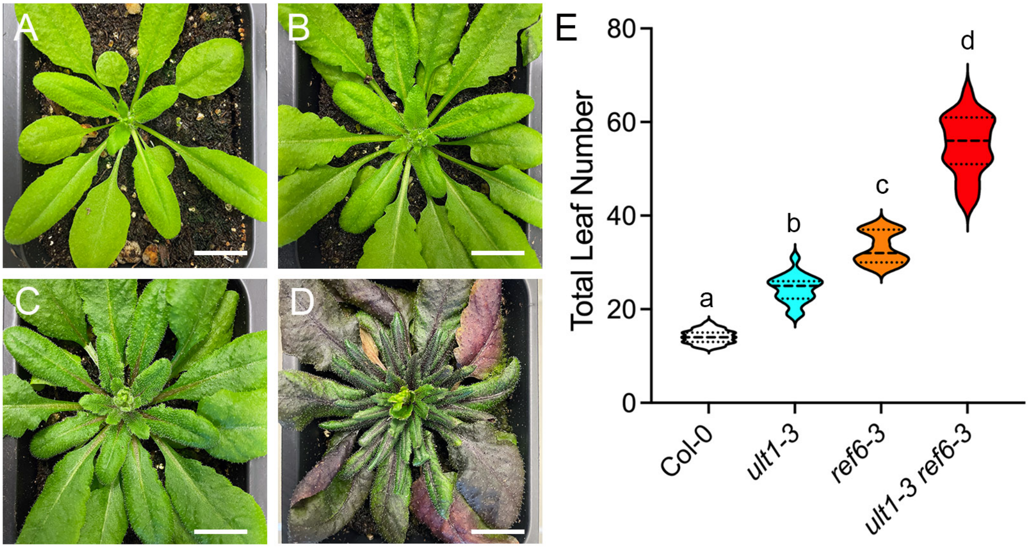
A
B
C
D
E
Total Leaf Number
80
60
40
20
0
a
b
c
d
Col-0
ult1-3
ref6-3
ult1-3 ref6-3

**Supplemental Figure 7.** *REF6* mutations enhance the *ult1* flowering time phenotype. **(A-D)** Top-down views of **(A)** wild-type Col-0, **(B)** *ult1-3,* **(C)** *ref6-3,* and **(D)** *ult1-3 ref6-3* plants grown under LD conditions. **(E)** Mean total leaf number. Lower case letters indicate statistically significant differences (P < 0.001). n > 20. Scale bar, 1 cm.

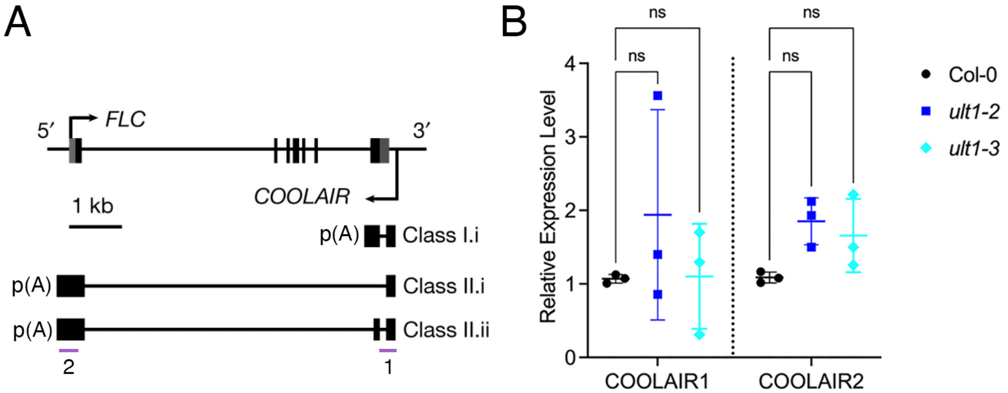
A
5′
FLC
3′
COOLAIR
1 kb
p(A)
Class I.i
p(A)
Class II.i
p(A)
Class II.ii
2
1
B
ns
ns
ns
ns
Relative Expression Level
4
3
2
1
0
COOLAIR1
COOLAIR2
Col-0
ult1-2
ult1-3

**Supplemental Figure 8.** ULT1 does not affect *COOLAIR* transcript levels. **(A)** Schematic of the *FLC* genomic region and the two classes of *COOLAIR* antisense transcripts, Class I and Class II (consisting of Class II.i and Class II.ii transcripts). Arrows indicate the *FLC* and *COOLAIR* transcription start sites and black bars indicate exons. Purple bars indicate primer set 1, which detects both Class I and Class II transcripts, and primer set 2 that detects only Class II transcripts. **(B)** Expression levels of the *COOLAIR1* (Class I) and *COOLAIR2* (Class II) transcripts in 12 DAG Col-0, *ult1-2* and *ult1-3* seedlings grown under LD conditions. Values indicate mean ± standard deviation for three biological replicates. ns = not significant.

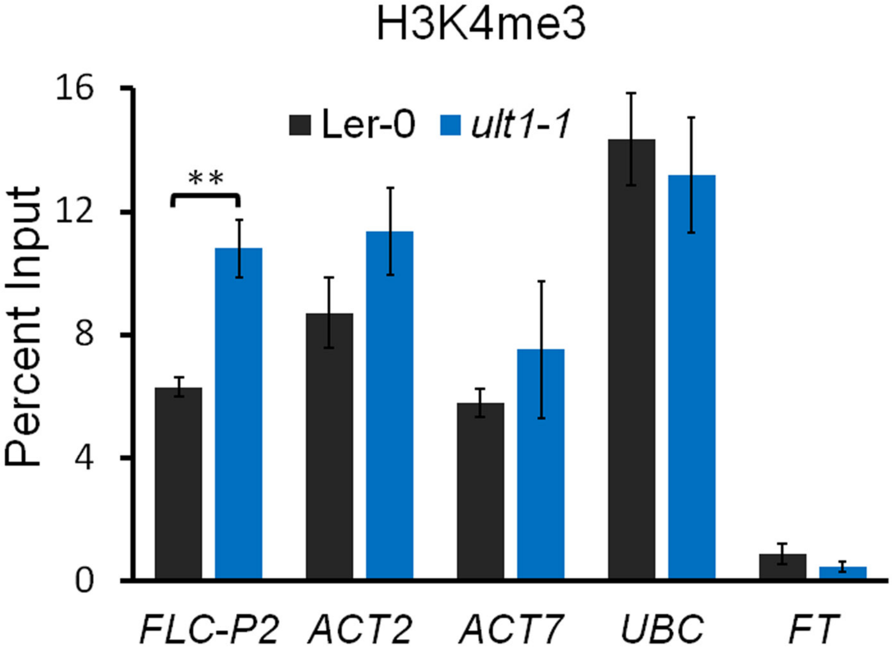

H3K4me3
Ler-0
ult1-1
**
Percent Input
16
12
8
4
0
FLC-P2
ACT2
ACT7
UBC
FT

**Supplemental Figure 9.** ULT1 modulates H3K4me3 levels at *FLC* but not at other loci. H3K4me3 levels at the *FLC-P2, ACT2, ACT7, UBC* and *FT* loci in 14 DAG L*er-0* and *ult1-3* plants. Values indicate mean ± standard error for three biological replicates. ** P < 0.01.

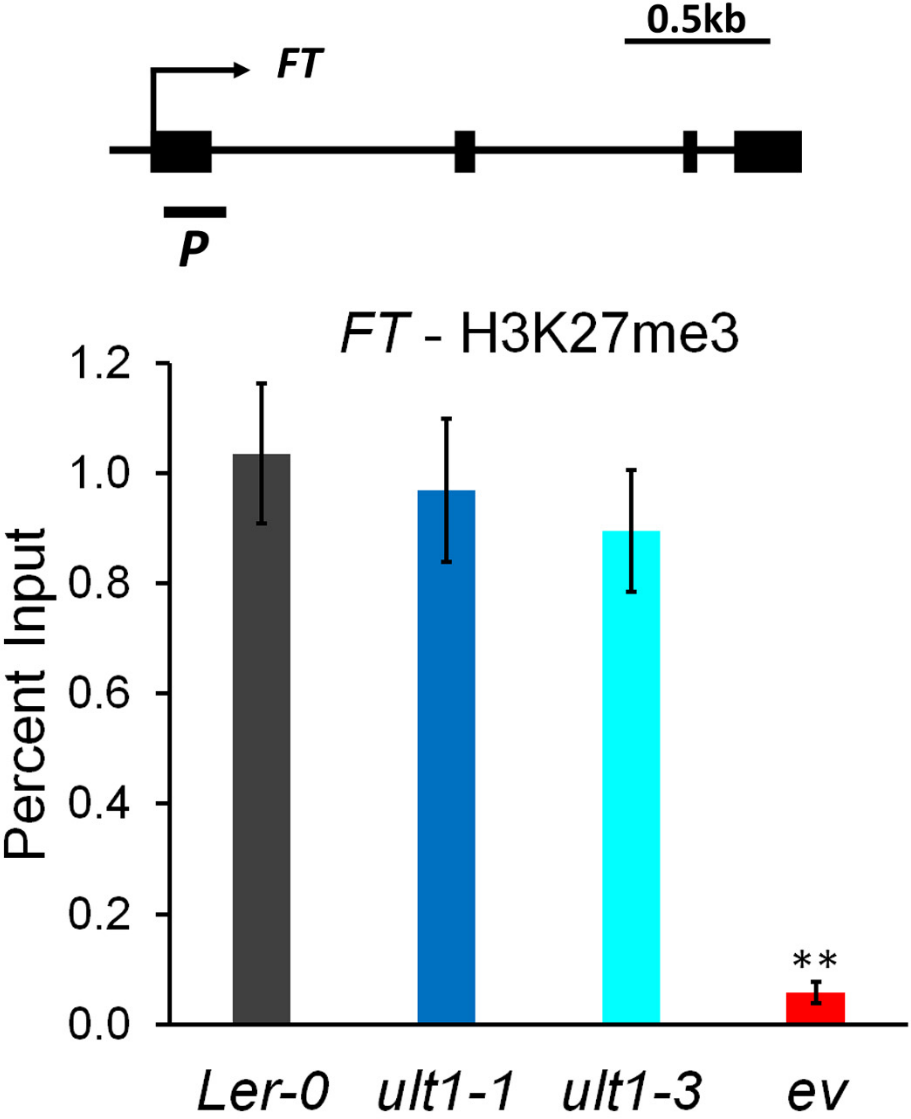

0.5kb
FT
P
FT - H3K27me3
Percent Input
1.2
1.0
0.8
0.6
0.4
0.2
0.0
**
Ler-0
ult1-1
ult1-3
ev

**Supplemental Figure 10.** ULT1 does not affect H3K27me3 levels at the *FT* locus. **(A)** Schematic of the *FT* genomic region. Large boxes show exons and the small boxes show UTRs. Arrow indicates the transcription start site (TSS). Bold dash indicates the position of the primers used in the ChIP-qPCR analysis. **(B)** H3K27me3 levels at the *FT* first exon in 14 DAG L*er-0*, *ult1-3* and *emf2-10 vrn2-1* plants. Values indicate mean ± standard error for three biological replicates. ** $P < 0.01$.

**Supplemental Table 1. List of prim**

**Genotyping**

**Primer Name**

flc-3 P2
flc-3 M2
ref6-3-P
ref6-3-M
ULT1F (ult1-3)
ULT1R (ult1-3)
ult2-4 M2
ult2-4 P2
LBb1.3
pDAP101

**Constructs**

**Primer Name**

ULT1_Y2H_F
ULT1_Y2H_R
CLF_Y2H_F
CLF_Y2H_R
SWN_Y2H_F
SWN_Y2H_R
EMF1_Y2H_F
EMF1_Y2H_R
ULT1-cLUC-F
ULT1_cLUC_R
ULT1_nLUC_F
ULT1_nLUC_R
SWN_nLUC_F
SWN_nLUC_R
SWN_cLUC_F
SWN_cLUC_R
FVE_cLUC_F
FVE_cLUC_R
CLF_nLUC_F
CLF_nLUC_R
CLF_cLUC_F
CLF_cLUC_R
ULT1_CDS_greengate_F
ULT1_CDS_greengate_R
CLF_CDS_greengate_F
CLF_CDS_greengate_R
SWN_CDS_greengate_F
SWN_CDS_greengate_R

**RT-qPCR**

| Primer Name |
|---|
| qRT_CO_F |
| qRT_CO_R |
| qRT_FLC_F |
| qRT_FLC_R |
| qRT_MAF4_F |
| qRT_MAF4_R |
| qRT_MAF5_F |
| qRT_MAF5_R |
| qRT_SVP_F |
| qRT_SVP_R |
| qRT_FT_F |
| qRT_FT_R |
| qRT_SOC1_F |
| qRT_SOC1_R |
| qRT_TSF_F |
| qRT_TSF_R |
| qRT_LFY_F |
| qRT_LFY_R |
| qRT_AP1_F |
| qRT_AP1_R |
| qRT_Total COOLAIR_F |
| qRT_Total COOLAIR_R |
| qRT_Distal COOLAIR_F |
| qRT_Distal COOLAIR_R |
| qRT_Proximal COOLAIR_F |
| qRT_Proximal COOLAIR_R |
| qRT_eIF4A_F |
| qRT_eIF4A_R |

**ChIP-qPCR**

| Primer Name |
|---|
| qChIP_FLC_P1_F |
| qChIP_FLC_P1_R |
| qChIP_FLC_P2_F |
| qChIP_FLC_P2_R |
| qChIP_FLC_P3_F |
| qChIP_FLC_P3_R |
| qChIP_FLC_P4_F |
| qChIP_FLC_P4_R |
| ChIP-FLC-P4'-F1 (ULT1-myc ChIP) |
| ChIP-FLC-P4'-R1 (ULT1-myc ChIP) |
| qChIP_FLC_P5_F |

qChIP_FLC_P5_R
qChIP_FLC_P6_F
qChIP_FLC_P6_R
qChIP_FLC_P7_F
qChIP_FLC_P7_R
qChIP_FLC_P8_F
qChIP_FLC_P8_R

**ers used in this study.**

| Sequence (5' to 3') |
| --- |
| TTGCATCACTCTCGTTTACCC |
| CGGAGGAGAAGCTGTAGAGCT |
| CAGGGATTCGACAACAATGTT |
| CAGTTGCAACTCTGGAGAAGG |
| ATGGCGAACAATGAGGGAGAG |
| TCAAGCTTTGACATTGCTGG |
| CACATTCTCTCTCCATCTCCG |
| TGGAGAATAAGGACCAGATGC |
| ATTTTGCCGATTTCGGAAC |
| TAGCATCTGAATTTCATAACCAATCTCGATACAC |

| Sequence (5' to 3') |
| --- |
| ATATGAATTCATGGCGAACAATGAGGGAGA |
| ATATGGATCCTCAAGCTTTGACATTGCTGGTG |
| AAAAGGCCATGGAGGCCATGGCGTCAGAAGCTTCGCC |
| AAAACCCGGGCTAAGCAAGCTTCTTGGGTC |
| AAAAGGCCATGGAGGCCATGGTGACGGACGATAGCA |
| AAAACCCGGGATGAGATTGGTGCTTTCTGG |
| AAAAGGCCATGGAGGCCATGGGATCTTCCATCAAGATC |
| AAAACCCGGGGGCATTTTGTGTAGGTGGACA |
| ACGCGTCCCGGGGCGGTACCATGGCGAACAATGAGGGAGAGA |
| AGCTCTGCAGGTCGACTCAAGCTTTGACATTGCTGGTGAAGT |
| CGGGGGACGAGCTCGGTACCATGGCGAACAATGAGGGAGAGA |
| ACGAGATCTGGTCGACAGCTTTGACATTGCTGGTGAAG |
| AGAACACGGGGGACGATGGTGACGGACGATAGCAAC |
| GACGCGTTGTGGATCCATGAGATTGGTGCTTTCTGGCT |
| CGGTACCTCCGGATCCATGGTGACGGACGATAGCAAC |
| TACGAACGAAAGCTCATGAGATTGGTGCTTTCTGGCTCTACG |
| ACGCGTCCCGGGGCGGTACCATGGAGAGCGACGAAGCAG |
| AGCTCTGCAGGTCGACTTAAGGCTTGGAGGCACAAGTCATAAC |
| CGGGGGACGAGCTCGGTACCATGGCGTCAGAAGCTTCGC |
| ACGAGATCTGGTCGACAGCAAGCTTCTTGGGTCTACCA |
| ACGCGTCCCGGGGCGGTACCATGGCGTCAGAAGCTTCGC |
| AGCTCTGCAGGTCGACCTAAGCAAGCTTCTTGGGTCTACCA |
| AACAGGTCTCAGGCTCAATGGCGAACAATGAGGGAGAG |
| AACAGGTCTCACTGAAGCTTTGACATTGCTGGTGAAGTC |
| AACAGGTCTCAGGCTCCATGGCGTCAGAAGCTTCGC |
| AACAGGTCTCACTGAAGCAAGCTTCTTGGGTCTACCAAC |
| AACAGGTCTCAGGCTCCATGGTGACGGACGATAGCAACT |
| AACAGGTCTCACTGAATGAGATTGGTGCTTTCTGGCTCT |

**Sequence (5' to 3')**
GGGAGATAGAGTTGTTCCG
GGATGAAATGTATGCGTTATGG
CGGTCTCATCGAGAAAGCTC
CCACAAGCTTGCTATCCACA
GCTTCTCCTCAGGTGATAGCATG
CTGCTCTTCCAGGGACTTTAGAC
TGTGTCGGAAGAGTGAAGCCAT
CTGATGATCTTGGCCATGCTGT
CAAGGACTTGACATTGAAGAGCTTCA
CTGATCTCACTCATAATCTTGTCAC
CAAAGAAAGGAGAAGGAGATACAG
CTGTTTGCCTGCCAAGCTGTC
ACGAGAAGCTCTCTGAAAAGTGGG
CTTGGGCTACTCTCTTCATCACCT
ATATCTCCACTGGTTGGTGAC
TGCATAAACCGTTTGTCTTCC
TCTCCCAAGAAGGGTTATCTG
TCTTCATCTTTCCTTGACCTG
CTCTCGTTTCTCACCACAACTC
TAAACGGGTTCAAGAGTCAG
TGCATCGAGATCTTGAGTGTATGT
ACGTCCCTGTTGCAAAATAAGC
GCTTCTCCTCCGGCGATAAG
AGAAAAGTAAAAGAGCACAAAACAGAA
GACCTATGATTATCGTACAGATGGAGA
GGCTAATTAAGTAGTGGGAGAGTCA
TTCGCTCTTCTCTTTGCTCTC
GAACTCATCTTGTCCCTCAAGTA

**Sequence (5' to 3')**
TTCAAGTCGCCGGAGATACT
CGTGGCAATCTTGTCTTCAA
CGACAAGTCACCTTCTCCAAA
AGGGGGAACAAATGAAAACC
ATCACAAGACTAATGATTAATGTCTC
GTCAATAGCTGCACAATGTGG
GAGGCACCAAAGAAACAAGG
TCGCCCTTAATCTTATGATCG
AACGAATTTCTCTCCTTTTTATGGG
AACACAACGATGATAAGATTAAGGG
TTTATCTGTCTTAGTCGCTTCC

GGTTCCAATTTCCTATATTTAAACCC
CCGGTTGTTGGACATAACTAGG
CCAAACCCAGACTTAACCAGAC
GATTTCAACCGCCGATTTAAGGTG
CATCATGTGGGAGCAGAAGCTG
CCTACGGCTTATTTTGCAAC
ACCATACGCGTTTAGGATAAA

**Supplemental Table 1.** Primer sequences used in the experiments.
